# Complex Autoinflammatory Syndrome Unveils Fundamental Principles of *JAK1* Transcriptional and Biochemical Function

**DOI:** 10.1101/807669

**Authors:** Conor Gruber, Jorg Calis, Sofija Buta, Gilad Evrony, Jerome Martin, Skyler Uhl, Rachel Caron, Lauren Jarchin, David Dunkin, Robert Phelps, Bryn Webb, Jeffrey Saland, Miriam Merad, Jordan Orange, Emily Mace, Brad Rosenberg, Bruce Gelb, Dusan Bogunovic

## Abstract

Autoinflammatory disease can result from monogenic errors of immunity. We describe herein the first example of a patient with early-onset widespread autoinflammation resulting from a mosaic, heterozygous, gain-of-function mutation (S703I) in *JAK1*, encoding a kinase essential for signaling downstream of over twenty-five cytokines. By first-of-its-kind custom single-cell RNA sequencing, we examine mosaicism with single cell resolution. We uncover that *JAK1* transcription is predominantly restricted to a single allele across different immune cells, introducing the concept of a mutational “transcriptotype” that differs from the genotype. Functionally, the S703I mutation not only increased JAK1 kinase activity, but also resulted in transactivation of partnering JAKs, independently of its catalytic domain. Further, S703I JAK1 was not solely hypermorphic for cytokine signaling, but neomorphic as well, as it enabled downstream signaling cascades not canonically mediated by JAK1. Given these results, the patient was treated with tofacitinib, a JAK inhibitor, which led to rapid resolution of her clinical disease. Together, these findings represent an unprecedented degree of personalized medicine with the concurrent discovery of fundamental biological principles.

## Introduction

Monogenic disease mutations afford the opportunity to study the *bona fide* function of human genes *in vivo*, which have guided our understanding of biology and medicine for decades. Undiagnosed disease programs, by means of next-generation sequencing, have recently provided a means to identify, diagnose, and study these rare patients with unusual clinical presentations (Splinter *et al*., 2018; Lee *et al*., 2019; Yang *et al*., 2019). In turn, clinical management can, in some cases, be highly personalized.

To date, studies of rare immunologic diseases have identified germline gain-of-function (GoF) and loss-of-function (LoF) mutations throughout the JAK-STAT signaling axis (Macchi *et al*., 1995; Russell *et al*., 1995; Dupuis *et al*., 2001; Kofoed *et al*., 2003; Minegishi *et al*., 2006, 2007; Holland *et al*., 2007; Mead *et al*., 2012; Hambleton *et al*., 2013; Etheridge *et al*., 2014) the primary signal transduction pathway for cytokines. The Janus kinase (JAK) family contains four tyrosine kinases (JAK1, JAK2, JAK3, TYK2) constitutively associated with cytokine receptors. Upon cytokine binding, JAKs act in partnership to phosphorylate themselves, the receptors, and then STATs, which can then act directly as transcription factors or activate other signaling pathways further downstream (O’Shea *et al*., 2015). Uniquely, JAK1 is activated by a remarkably broad range of cytokines (γc-, gp130-, Interferon- and IL-10-family cytokines). It can phosphorylate any STAT protein (STAT1-6), and is universally expressed in all tissues (O’Shea *et al*., 2015). Through the formation of specific combinations of cytokine receptors, JAK partners, and STAT dimers, JAK1 orchestrates unique downstream signals for each cytokine.

The need to better understand JAK regulation has deepened with the expanding clinical use of JAK inhibitors (O’Shea and Gadina, 2019). The breadth of successfully treated inflammatory conditions signifies the central pathophysiological role of JAK hyperactivity across immune diseases. Yet, the complete list of disorders resulting from JAK-STAT dysregulation remains unknown. Further, it is unclear which specific JAK-mediated pathways drive disease, a key issue for the design of inhibitors with greater selectivity for individual members of the JAK family.

Herein, we identify a novel mutation (S703I) of *JAK1* in a patient with a severe, early-onset immunodysregulatory syndrome identified in our undiagnosed disease program. Using extensive next-generation genomic, molecular and multi-parametric immunological tools, we probe the effects of S703I *JAK1 in vitro* and *ex vivo* in order to investigate clinical dysfunction *in vivo*.

## Results

### Complex immunodysregulatory syndrome

We studied an 18 year-old female from a non-consanguineous family who was referred into our undiagnosed disease program with an early-onset, severe syndrome of immune dysregulation. Both her parents and an older brother were healthy (Figure 1A). In the patient, disease features arose prior to the age of 3 years, and included: diffuse, severe dermatitis; chronic unspecified enteritis and colitis with increased eosinophils; idiopathic membranous nephropathy (MN); fluctuating peripheral eosinophilia; asthma; food and environmental allergies; severely stunted growth with leg length discrepancy and poor weight gain (Figure S1A-E). Of note, serology was negative for known MN autoantibodies. Similarly, neither gastrointestinal nor dermatologic manifestations matched criteria for known diagnostic entities, despite multiple clinical evaluations and biopsies. However, the skin disease was reminiscent of atopic dermatitis, and the linear lesions were similar to those found in inflammatory linear verrucous epidermal nevus (ILVEN). Kidney transplantation was performed at age 11, but MN recurred within one year, followed by antibody-mediated rejection, requiring subsequent hemodialysis. The other inflammatory and autoimmune features were also largely resistant to therapeutic interventions (Supplemental Case Report).

**Figure 1.**
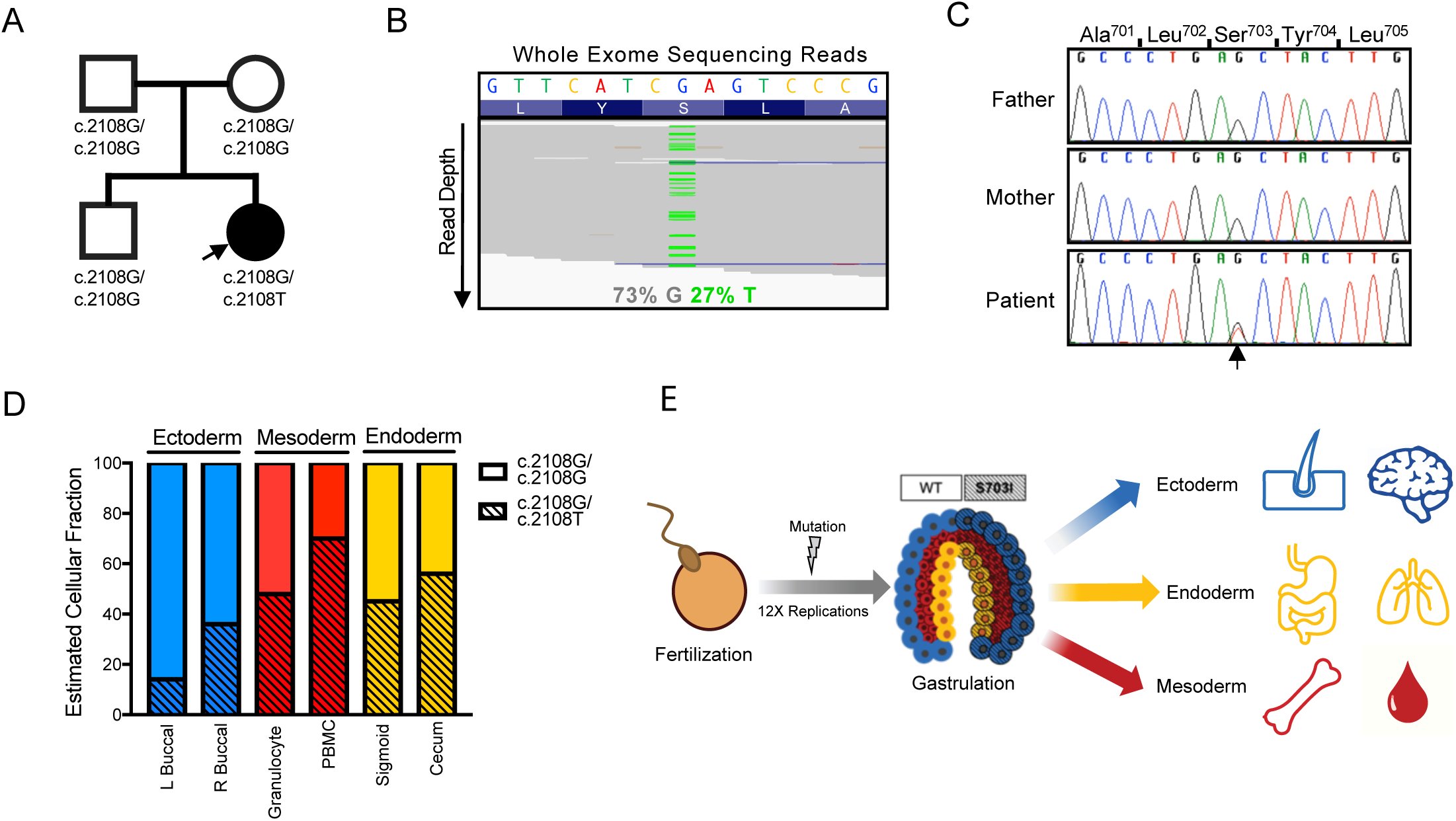
*De novo* mutation in *JAK1* identified in patient with immunodysregulatory syndrome. (A) Patient’s family pedigree. (B) Whole exome sequencing reads mapping to *JAK1* locus c.2108 with variant nucleotides displayed in green (C) Representative chromatograms from Sanger sequencing of peripheral blood DNA to confirm c.2108 G>T *JAK1* (D) Proportion of cells carrying heterozygous mutation, as estimated by digital droplet PCR with WT- and mutation-specific probes. DNA was obtained from bilateral cheek swabs, Ficoll-fractionated whole blood, and epithelial tissue isolated from a colonic biopsy (n=1). (E) Model for the development of the *de novo* mutation and its distribution into all three germ layers.

### Whole exome sequencing reveals a *de novo JAK1* mutation

Given the overall healthy state of the parents, and the early onset of disease in the patient, we hypothesized that either a recessive or a *de novo* genetic mutation was causative of the clinical syndrome. We performed whole-exome-sequencing on peripheral blood cells obtained from the patient and her parents. Subsequent variant analysis failed to produce any likely variants by a recessive model of inheritance (Table S1). Given the asymmetric manifestations of disease, including limb-length discrepancy and irregularly distributed dermatitis, we then considered the possibility of lower-read-frequency *de novo* mosaic mutations, which are typically excluded from common analysis pipelines. One candidate *de novo* variant, *JAK1* c2108G>T, which constituted 27% of reads mapping to the region, was identified (Figure 1A-B). The presence of the c2108G>T variant was confirmed by Sanger sequencing (Figure 1C), and this variant was absent from all publicly available genome sequences. This mutation results in substitution of serine to isoleucine at position 703 (S703I) in a highly conserved region (Figure S1F) and is predicted to be highly damaging (CADD score of 27.6). We then investigated the presence of c.2108G>T in non-hematopoietic tissues. We performed digital droplet PCR (ddPCR) with mutation-specific probes to estimate the fraction of cells carrying the mutation in different tissues. We identified the mutation at various frequencies in DNA from buccal swabs, granulocytes, PBMCs, and endoscopic biopsy samples fractionated into epithelia and associated immune cells (Figure 1D and S1G). These tissues represent all three germ layers, signifying that the mutation must have arisen in the first ∼12 cell divisions between fertilization and gastrulation (Figure 1E)(Moore, Persaud and Torchia, 2015).

### Allele characterization indicates that S703I confers a gain-of-function on JAK1

The S703I mutation localizes to the pseudokinase domain of JAK1, a putative regulatory domain (Figure 2A). Although S703I is positioned adjacent to the two germline *JAK1* mutations identified to date, these other mutations diverged in their downstream consequences, making functional predictions for S703I difficult. To assess the possible pathogenicity of the mutation and its impact on JAK1 function, we transduced WT *JAK1*, S703I *JAK1* and empty vector lentiviruses into U4C (*JAK1-/-*) cells. Transduction with S703I *JAK1*, but not WT *JAK1* or *Luciferase*, led to basal phosphorylation of STAT proteins and active target gene transcription in the absence of cytokine stimulation (Figure 2B-E). S703I-transduced cells hyper-responded to IFNα, in terms of both the proximal phosphorylation of STAT1 and STAT2 and the induction of interferon-stimulated genes (ISGs) (Figure 2B-C). Similarly, these cells hyper-phosphorylated downstream STATs in response to IFNγ or IL-6 (Figure 2D-E and S2A).

**Figure 2.**
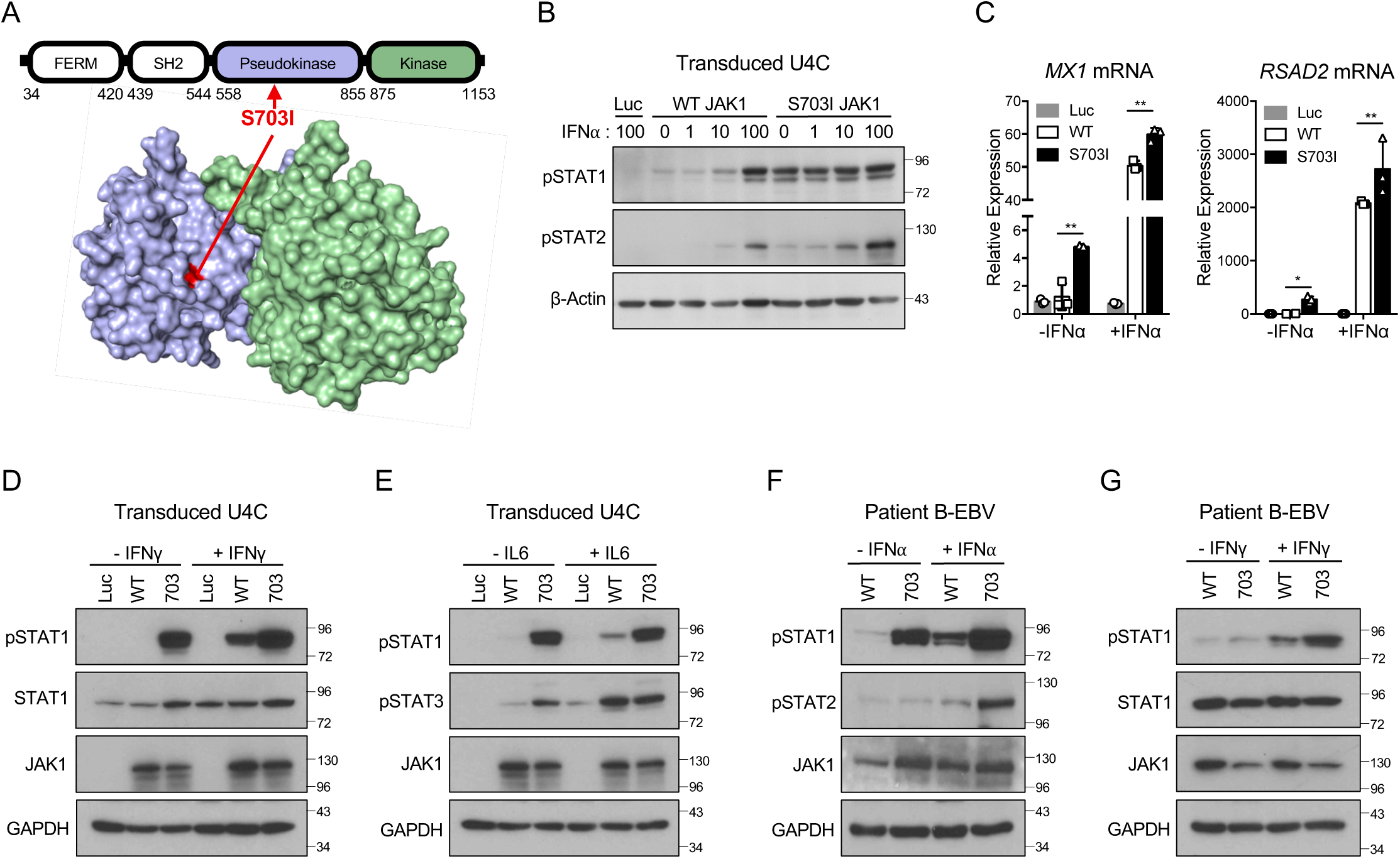
S703I JAK1 confers constitutively active and hyper-responsive STAT signaling. (A) Localization of S703I (red) mutation to the pseudokinase domain (blue) of JAK1, as represented in the linear sequence and predicted structure modeled on TYK2 (B) Western blotting for STAT phosphorylation in U4C Cells (*JAK1-*/-) reconstituted with lentiviruses for vector control (Luciferase), WT *JAK1*, and S703I *JAK1* and stimulation for 15 minutes with indicated doses of IFNα (IU/mL). (C) qPCR from transduced U4C cells for interferon-stimulated gene expression (*MX1* and *RSAD2*) at baseline and following 100 IU/mL of IFNα for 8 hours (n-3). Columns represent mean and error bars represent standard deviation. *P < 0.05. **P < 0.01, ***P < 0.001, two-tailed Student’s t test with Welch’s correction. (D) Stimulation of transduced U4C cells with IFNγ (0.1 ng/mL) or (E) IL6 (25 ng/mL) for 15 minutes. (F) Derivation of BEBV cells and isolation of JAK1 WT/WT and JAK1 S703I/WT cells from patient blood, followed by stimulation with IFNα (100 IU/mL) or (G) IFNγ (0.1 ng/mL) for 15 minutes.

For direct confirmation of the pathogenicity of the mutation in cells from the patient, we derived an EBV-immortalized B-cell (B-EBV) line from the patient’s PBMCs. Given the mosaicism for *JAK1* in the patient’s cells, individual lines were cloned from single cells to derive purely WT or S703I heterozygous mutant cells (Figure S2B). A comparison of STAT phosphorylation in patient WT and mutant B-EBV cells supported the gain-of-function role of S703I JAK1, both at baseline and in response to cytokine (Figure 2F-G). The isogenic control derived from the same patient definitively identified S703I JAK1 as the probable pathogenic mutation in the patient’s genome. Together, these results indicate that S703I is gain-of-function for basal and cytokine-induced STAT signaling.

### S703I JAK1 trans-activates partnering JAKs independently of its own kinase activity

To dissect the mechanisms underlying upregulated STAT signaling, we assessed the impact of S703I on JAK auto-phosphorylation. Consistent with the increase in STAT phosphorylation, S703I JAK1 was itself hyper-phosphorylated (Figure 3A). Interestingly, both JAK2 and TYK2 phosphorylation were also upregulated (Figure 3B-C S2C), suggesting that the interacting JAK partners may also play a role in the gain-of-function. We hypothesized that the mutant JAK1 pseudokinase domain trans-activates the kinase activity of JAK2 and TYK2. To investigate this mechanism, we mutated the ATP-binding site (K908A) of JAK1 to render it catalytically inactive. This well-characterized mutation retains the signaling capability of the receptor complex, making it possible to study signaling by the partnering JAKs in isolation (Li *et al*., 2013; Eletto *et al*., 2016). As expected, STAT phosphorylation levels were largely reduced in the absence of JAK1 activity (Figure 3D-E). Yet, remarkably, following inactivation of the kinase domain of S703I JAK1 (S703I/K908A), an aberrant increase in STAT phosphorylation relative to kinase-inactivated JAK1 without the S703I mutation was observed upon cytokine stimulation (Figure 3D-E). This result indicates that S703I JAK1 trans-activates JAK2 and TYK2, revealing that pseudokinase domains can regulate partnering JAKs in *trans*, in addition to traditionally understood *cis*-regulation (Babon *et al*., 2014). This novel mechanism of JAK-STAT signaling regulation, revealed by the rare disease mutation of a single patient, highlights both the potential of rare disease research to discovery of novel physiological mechanisms and the unexpected complexity with which mutations of a kinase gene can enhance signaling independently of their own activity.

**Figure 3.**
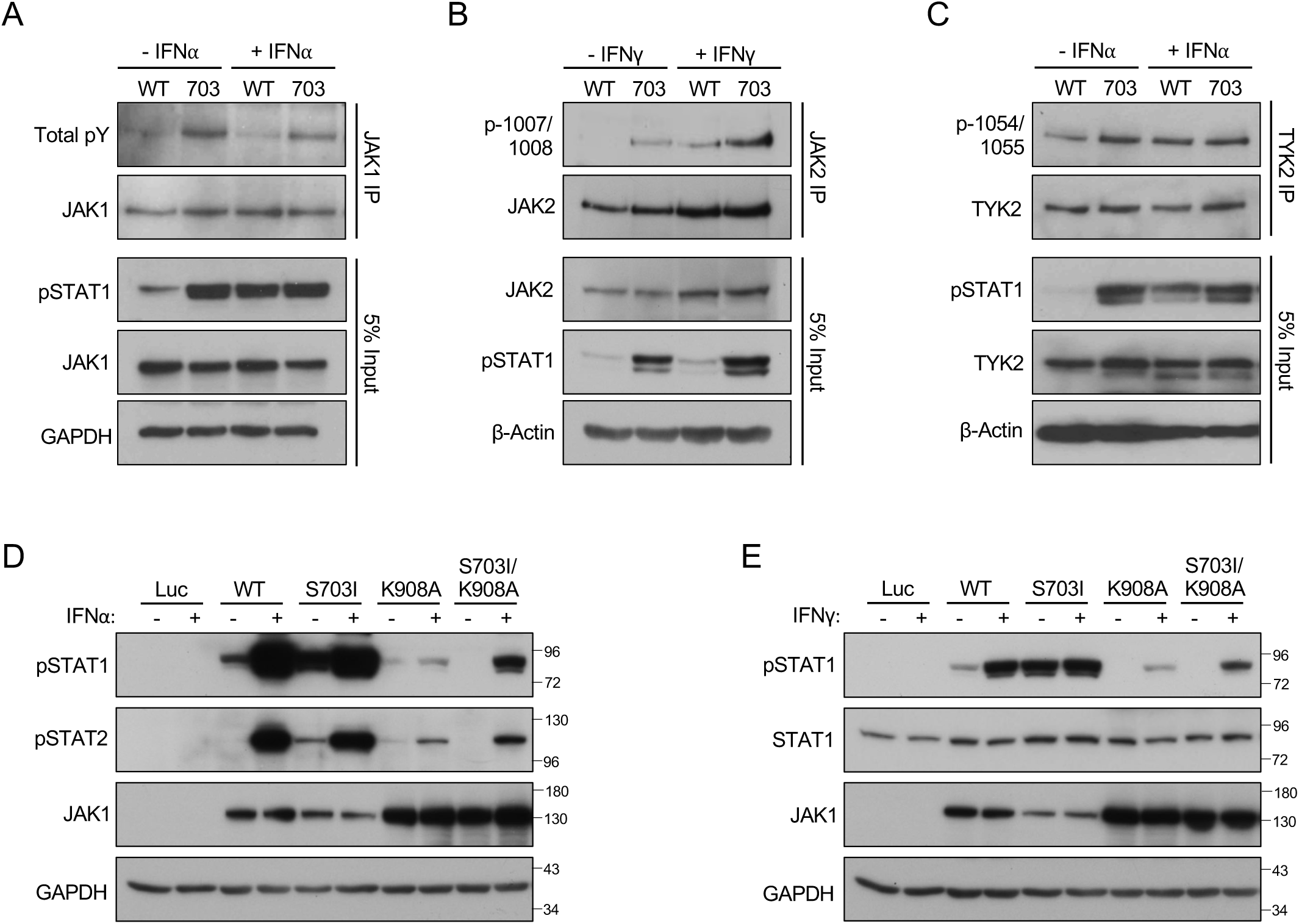
S703I JAK1 mediates hyperactive STAT phosphorylation by trans-activation of partnering JAKs. (A) Immunoprecipitation of JAK1 from *JAK1*-transduced U4C cells and western blotting for total phosphorylation (4G10) after stimulation with IFNα (100 IU/mL) for 15 minutes. (B) Immunoprecipitation of JAK2 from *JAK1*-transduced U4C cells and western blotting for phosphorylation at the activation loop after stimulation with IFNα (100 IU/mL) for 15 minutes. (C) Immunoprecipitation of TYK2 from patient-derived B-EBV cells and western blotting for phosphorylation at the activation loop after stimulation with IFNγ (0.1 ng/mL) for 15 minutes. (D). Transduction of U4C cells with catalytically inactivated *JAK1* (K908A), WT *JAK1*, S703I *JAK1*, or double mutant *JAK1* (K908A S703I), followed by stimulation with IFNα (1000 IU/mL) or (E) IFNγ (1.0 ng/mL).

### *Ex vivo* analysis reveals an expansion of CD56^hi^ NK cells and a cell-intrinsic gain-of-function

To more robustly investigate the consequences of S703I on immune cells, we performed mass cytometry (CyTOF) immunophenotyping on whole blood from the patient. Surprisingly, despite the central role of JAK1 in immune cell differentiation and proliferation, the patient’s immune cell distribution was largely within the normal range, barring a few exceptions (Figure 4A, Table S2). B cells, for instance, trended toward lower frequencies than in healthy donors. Most strikingly though, natural killer (NK) cells exhibited a dramatic expansion of CD56^hi^ cells (>12-fold over healthy controls) (Figure 4B, Figure S3A-B). The CD56^hi^ subset is the minority population in healthy individuals in whom about 90% of peripheral blood NK cells are CD56^lo^ (Poli *et al*., 2009). These subsets are broadly defined by their surface density of CD56 but have unique phenotypic and functional properties. Namely, the CD56^hi^ subset is less mature but rapidly produces cytokines, whereas the CD56^lo^ subset expresses perforin and granzyme at baseline and mediates cytotoxicity through contact-dependent target cell lysis (Poli *et al*., 2009). Extensive phenotyping of the patient’s NK cells demonstrated that the expanded population retained many prototypical immature features defining them as *bona fide* “CD56^bright^” NK cells (Figure S3C-E). The CD56^lo^ NK cell population was present and similarly expressed canonical defining markers, but it had not undergone such expansion.

**Figure 4.**
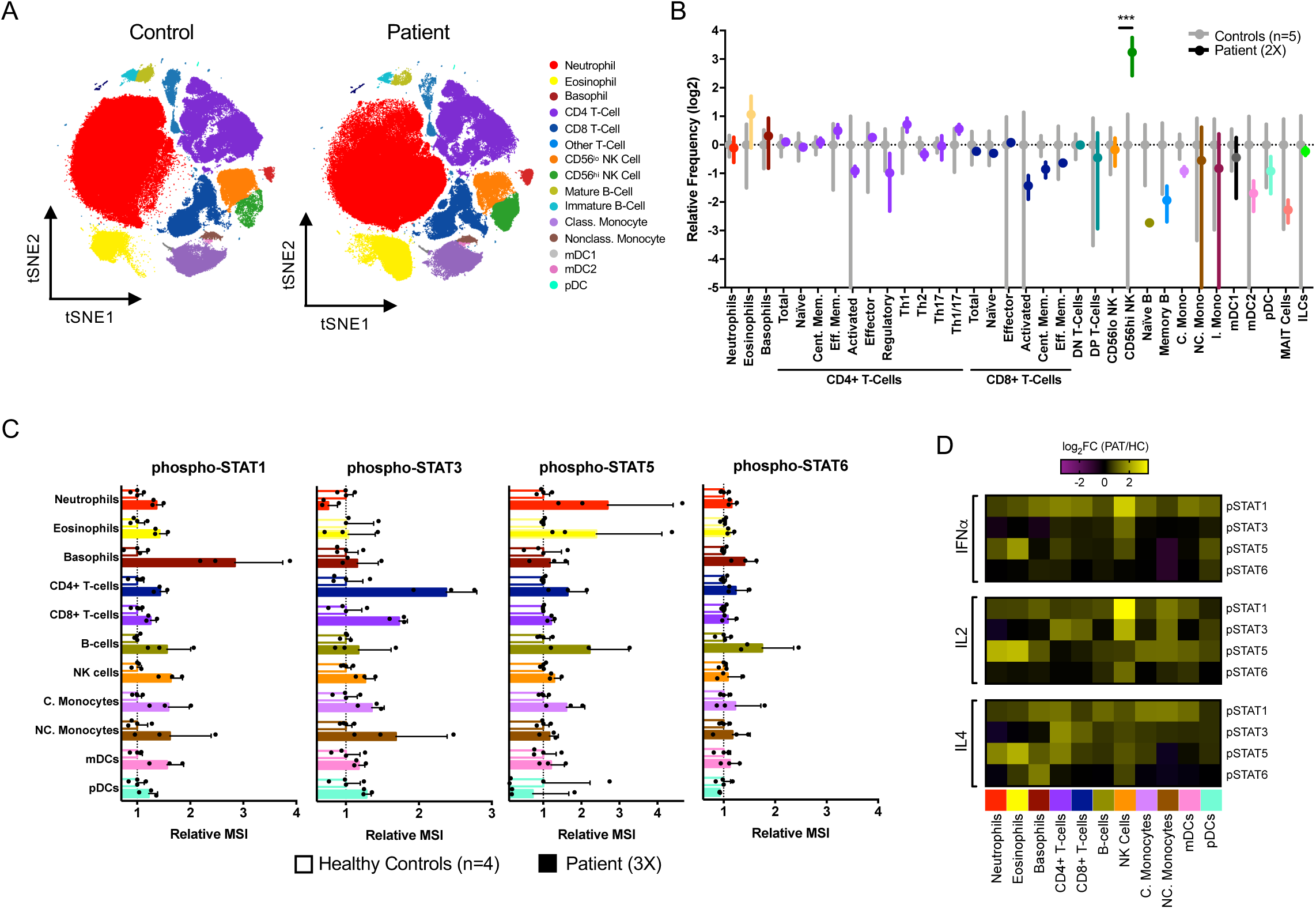
CyTOF analysis reveals cytokine-, STAT- and cell-type-specific gain-of-function. (A) Representative tSNE plots generated from immunophenotyping CyTOF data of whole blood. (B) Manually gated CyTOF populations from whole blood of five separate healthy controls and the patient on two separate occasions (2X) were quantified as percent of single cells and expressed as relative frequency (patient/controls). Grey bars indicating mean with standard deviation of healthy donors, and colored bars indicating mean with standard deviation of patient. Multiple t-tests performed correcting for multiple comparisons using the Holm-Sidak method. ***P < 0.001. (C) Relative MSI (patient/control) of phospho-STAT staining from intracellular phospho-CyTOF of whole blood from four healthy donors (n=4) and the patient on three separate occasions (3X). (D) *Ex vivo* stimulation with IFNα (100 IU/mL), IL-2 (50 ng/mL) and IL-4 (50 ng/mL) for 15 minutes. Color intensity indicates the log2 fold-change in MSI between patient and healthy control (n=1).

We then performed phospho-CyTOF to analyze the phosphorylation of all STATs downstream from JAK1 in all major immune cells of whole blood. Given the ubiquitous expression and diverse signaling capabilities of JAK1, as well as the basal activity of S703I observed *in vitro*, we hypothesized that all STATs within all immune cells would be hyper-phosphorylated at baseline. Indeed, heightened STAT phosphorylation was observed, but not universally. Certain immune subsets, but not others, exhibited high basal phosphorylation of specific STATs (Figure 4C). For example, granulocytes from the patient displayed baseline STAT1 phosphorylation, but not STAT3 phosphorylation, whereas T cells displayed basal STAT3 phosphorylation but not STAT1 phosphorylation. STAT6 phosphorylation, however, was not significantly upregulated in any immune subset except B cells.

To evaluate the functional consequences of the observed basal phosphorylation, we assessed the expression of downstream genes from bulk PBMCs. We detected elevated expression of many downstream pSTAT target genes, including *IFIT1* and *SIGLEC1* (Figure S4A). In addition, we tested non-hematopoietic tissues for baseline STAT phosphorylation by immunohistochemistry. In skin, gastrointestinal and renal biopsies we detected phosphorylated STAT1 and STAT3 (Figure S4B). This basal phosphorylation was observed in the apparent absence of any overt increase in circulating JAK-STAT cytokine levels (Figure S4C), suggesting that intrinsic S703I JAK1 activity drives this process, consistent with our *in vitro* results (Figure 2B-F).

### Ex vivo cytokine stimulation generates novel pathways in patient cells

Whole blood was then stimulated *ex vivo* with a series of cytokines that engage JAK1 with various cytokine receptors, JAK partners and downstream STAT targets. In response to IFNα or IL-2, patient leukocytes hyper-phosphorylated STAT1 and STAT3, or STAT5, respectively (Figure 4D, represented differently in Figure S5A-C). Of note, NK cells exhibited the strongest response to IFNα and IL-2. By contrast, STAT6 phosphorylation in response to IL-4 was similar to that in healthy control cells, as seen in the baseline STAT6 data (Figure 4D, S5C). The same differential response was noted in patient B-EBV cells (Figure S5D-E). Interestingly, CyTOF analysis revealed that both IL-4 and IL-2 induced the phosphorylation of STAT1 in patient cells, contrasting with the canonical signaling cascade induced by these cytokines (Fig 4D). This non-canonical response suggests that S703I confers promiscuity onto JAK1, allowing it to transverse traditional signaling axes and establish novel pathways. Together, these results indicate that S703I JAK1 is a gain-of-function mutation *ex vivo*, displaying both unexpected pathway promiscuity and cell-type specificity in the activation of STAT signaling.

### Custom scRNAseq maps cellular distribution and transcriptomic signatures of S703I *JAK1*

To follow up on these striking differences across the patient’s cell types, we performed single cell RNA sequencing of patient PBMCs. We aimed to specifically map and evaluate the impact of S703I JAK1 across immune cell subsets by relating single cell-resolution gene expression patterns to cell type and, given the mosaicism, *JAK1* genotype. However, sequence data from droplet-based scRNA-Seq platforms are typically restricted to 5’ or 3’ transcript regions and therefore do not include coverage of the S703I site (c.2108) in *JAK1* mRNA (Figure S6A). Therefore, using custom barcode microbeads and a modified library preparation procedure, we adapted the inDrop scRNA-Seq methodology (Klein *et al*., 2015; Zilionis *et al*., 2017) to target exon 16 (containing the S703I site) of JAK1 in addition to standard mRNA 3’ regions (Figure S6B, details in *Methods*). We used *JAK1*-targeted inDrop scRNA-Seq to analyze patient PBMCs, and concurrently processed the same sample on the 10X Genomics Chromium platform (Figure S6B). Clustering and manual annotation of cell types based on gene expression patterns distinguished the expected PBMC populations, with relative frequencies consistent with the corresponding CyTOF analyses (Figure 5A, S6C). Again, we observed a relative increase in CD56^hi^ NK cell frequency. Using sequence data from *JAK1*-targeted libraries, we assigned putative per cell *JAK1* genotypes (based on transcript sequences) to a subset of PBMCs for which transcript and read depths were sufficient in the *JAK1*-targeted inDrop dataset (Figure S6D-E). Within this subset, we found that the mutant allele was not evenly distributed across immune cell-type clusters (Figure 5B-C). For example, B cells and monocytes mostly carried WT transcript, whereas 69% of CD56^hi^ NK cells contained JAK1 mRNA with the S703I mutation. This variable distribution may reflect differences in the intrinsic tolerance to the mutation in these cell types, which is consistent with the expansion of CD56^hi^ NK cell subset and the higher levels of *JAK1* expression in NK cells than in other cell types (Figure S6F).

**Figure 5.**
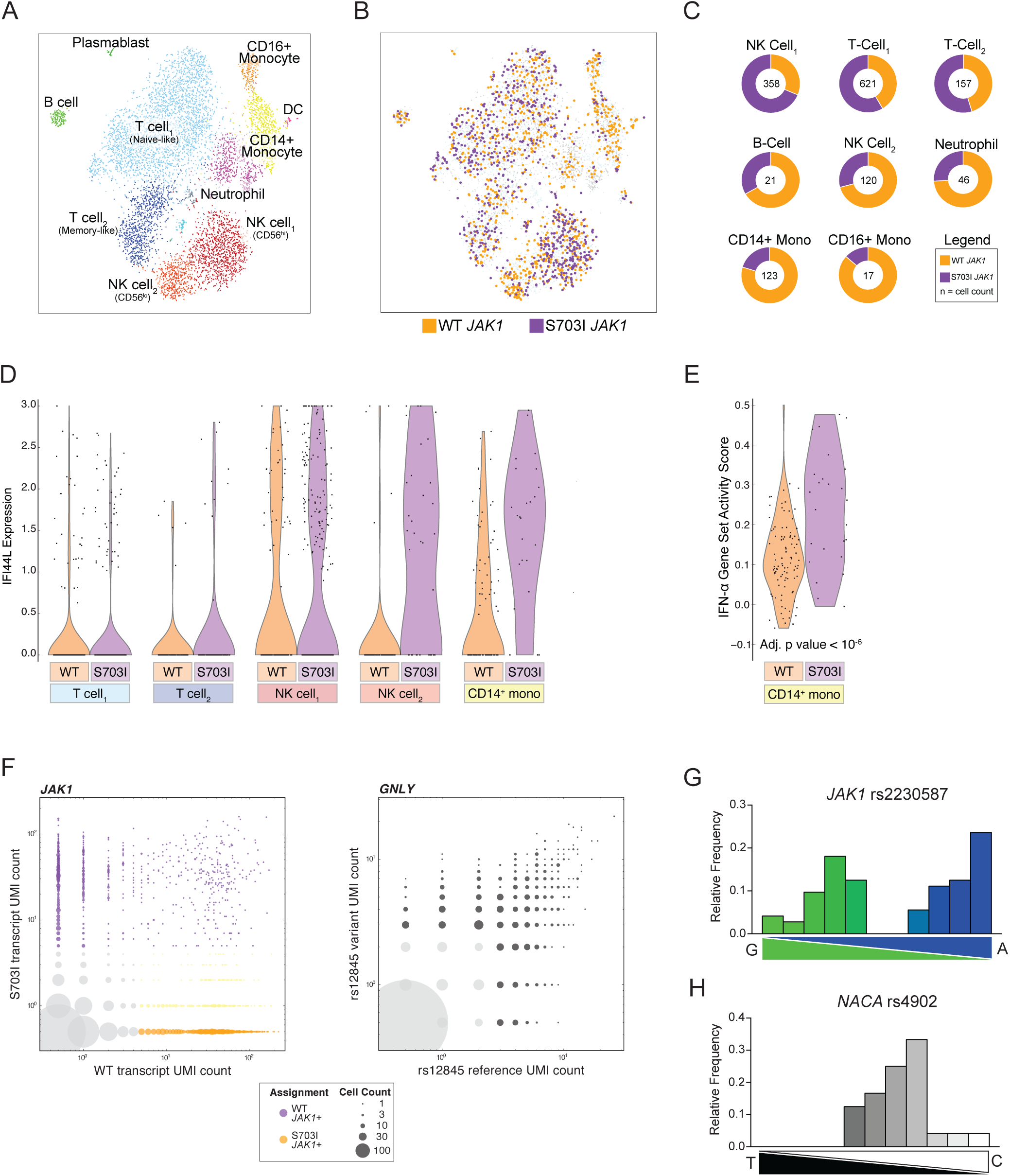
Custom single-cell-RNA sequencing maps JAK1 allele distribution, transcriptomic impact and expression patterns. (A) tSNE plots and cell type assignments from single cell RNA sequencing of patient PBMCs with an inDrop platform adapted to target the mutant JAK1 transcript. n=4763 cells. (B) tSNE plots representing the subset of cells with sufficient JAK1 counts to be assigned putative JAK1 genotypes (based on transcript sequences). Cells in which any mutant transcript was detected above empirically-determined thresholds were assigned ‘S703I JAK1’ (purple) while cells with only WT transcript detected were assigned ‘WT JAK1’ (orange). (C) Doughnut charts quantifying allele distribution in cells meeting genotyping criteria (cell count in the center), as in panel B. (D) Expression of the interferon stimulated gene IFI44L, a statistically significant differentially expressed gene in the comparison of WT JAK1 and S703I JAK1 genotyped cells. (E) Gene set scores for IFNα signaling in CD14+ monocytes (F) Number of unique transcripts detected per cell for the WT or S703I JAK1 allele (left) or a control variant GNLY rs12845 (right). Bubble size indicates number of cells. Color coding indicates cells containing: S703I JAK1 (purple), WT JAK1 without S703I JAK1 (orange), WT JAK1 with S703I JAK1 detected below threshold (yellow), or insufficient transcripts counts (gray). (G) Transcript genotyping of JAK1 rs2230587 from healthy control PBMCs (n=96) by single cell qPCR with allele-specific probes. Histogram represents relative frequency of cells expressing binned allele ratios as quantified by oligonucleotide standards. (I) Single cell qPCR transcript genotyping of control gene NACA (rs4902).

We next performed differential gene expression analysis, comparing patient cells expressing the WT and mutant alleles of *JAK1*. Although this analysis was constrained by a limited number of cells for which transcript-level genotype information was available, the intrasample comparison can be conducted on an inherently isogenic background with identical exposure history. We detected statistically significant differences in ISG expression for *IFI44L* and for the entire ISG gene set between WT and mutant monocytes (Figure 5D-E). These data suggest that tolerance to S703I may differ between cell types and confirm the cell-intrinsic nature of the gain-of-function described above.

### Biased expression of *JAK1* alleles

Given the mosaicism of a heterozygous mutation, we expected to observe some cells (i.e. homozygous WT *JAK1*) containing only WT *JAK1* transcripts and others (i.e. heterozygous S703I *JAK1*) containing both WT and S703I *JAK1* transcripts. Surprisingly, we found that expression of the two alleles seemed almost mutually exclusive, as very few cells expressed both transcripts, as opposed to the ∼50% expected given our genomic estimates (Figure 5F left panel, Figure 1D). By contrast, other genes from our dataset containing heterozygous variants exhibited the expected biallelic distribution (Figure 5F right panel). This result suggests that *JAK1* may be subject to monoallelic bias, a pattern that has only recently been recognized in a fraction of the autosomal transcriptome (Gimelbrant *et al*., 2007; Jeffries *et al*., 2012; Deng *et al*., 2014; Borel *et al*., 2015). We further tested this hypothesis of biased allele expression by sorting single cells from healthy donor PBMCs heterozygous for a synonymous SNP (rs2230587) of *JAK1*, adjacent to S703I. qPCR of isolated RNA with allele-specific probes revealed that relative expression levels of the two alleles were not normally distributed, but rather biased to one allele or the other (Figure 5G), unlike in a control gene (Figure 5H). Query of publicly available murine expression data revealed a similar allele restriction for *JAK1*, which, at least in mice, remains fixed over time (Savova, Patsenker and Gimelbrant, 2016). Whether the observed phenomenon in this patient represents transcriptional bursting or mitotically stable monoallelic expression, and the potential impact on immune dysfunction, remains to be fully determined. In either sense, these data, perhaps, indicate departure from the classic genetic interpretation of heterozygosity and allow for a shift in understanding of genetic penetrance of disease.

### Clinical and biological rescue with tofacitinib

Having identified JAK1 hyperactivity as the putative driver of clinical disease in the patient, we then considered the use of JAK inhibitors for the clinical treatment of this patient. We first compared the ability of the two FDA-approved inhibitors available at the time to reduce basal STAT phosphorylation in S703I-transduced U4C cells. Despite its lower relative potency against JAK1, tofacitinib inhibited STAT phosphorylation in a comparable dose response to ruxolitinib (Figure 6A), again reflecting the independence of JAK1 catalysis and STAT hyper-phosphorylation. Similar results were obtained with patient-derived B-EBV cells, which, unlike U4Cs, express JAK3 (Figure 6B). Next, we treated patient blood *ex vivo* with the two compounds at equimolar doses that mimic physiological dosing (Chen *et al*., 2014; Krishnaswami *et al*., 2014; Lamba *et al*., 2016), and we further assessed the inhibition of IFNα stimulation. Analysis of phospho-STAT inhibition across whole blood immune subsets by phospho-CyTOF revealed that tofacitinib attenuated the response more potently than ruxolitinib in nearly all cell types (Figure 6C).

**Figure 6.**
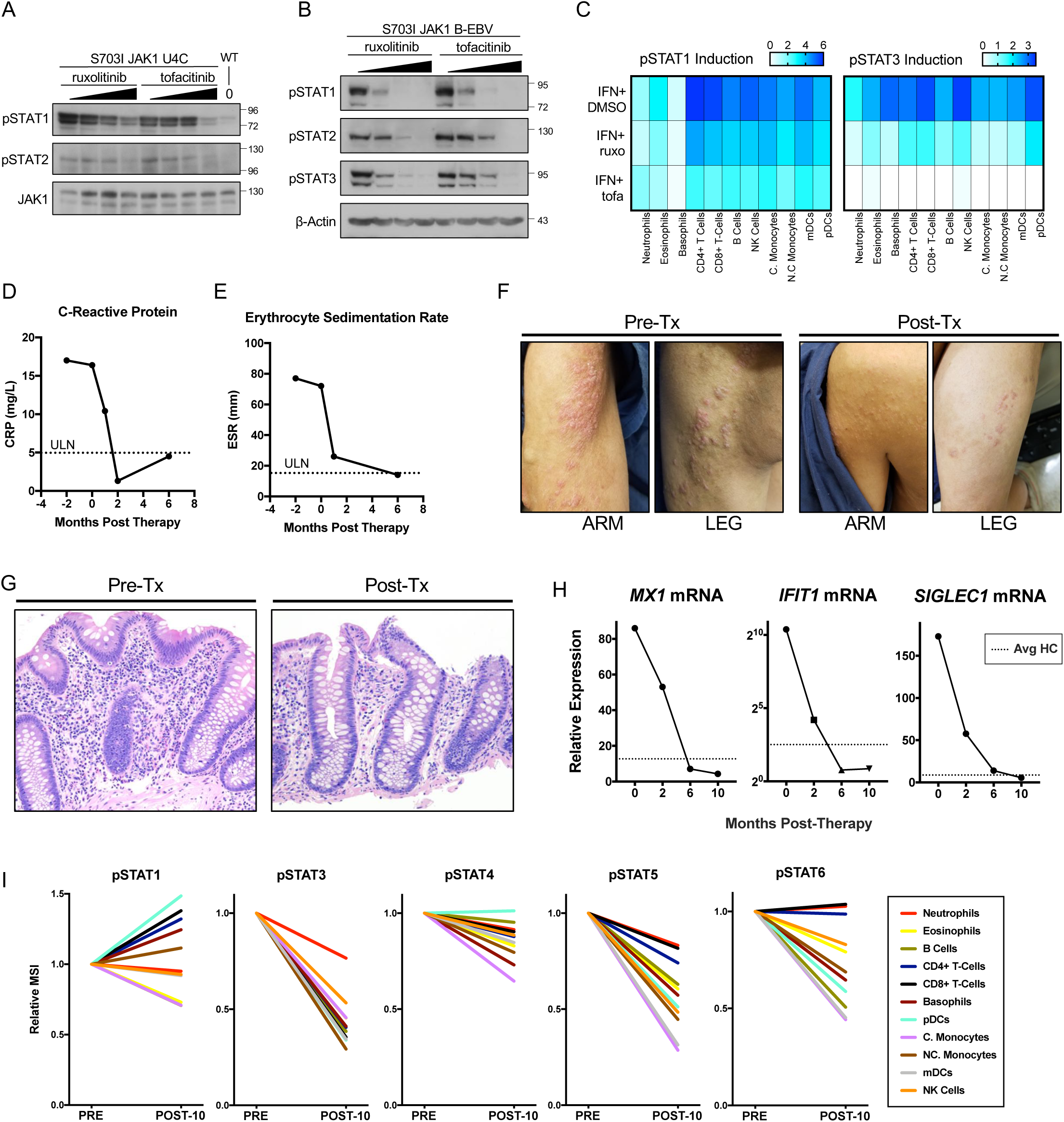
Treatment with tofacitinib rescues STAT hyperactivity and resolves clinical disease. (A) *In vitro* assessment of JAK-inhibitor efficacy after 4-hour drug treatment for resolution of basal STAT phosphorylation in transduced U4Cs. (B) Similar analysis in patient-derived B-EBVs. (C) *Ex vivo* inhibition with equimolar doses (500 nM) of ruxolitinib and tofacitinib for 4 hours followed by IFNα stimulation (1000 IU/mL) (n=1). (D & E) Acute phase reactants from peripheral blood. (F) Gross appearance of skin before and 5-months after tofacitinib initiation. (G) Colonic biopsies before and 6-months post-therapy. (H) ISG expression in RNA from bulk PBMCs isolated throughout treatment (n=1). (I) Phospho-CyTOF analysis comparing relative changes in pSTAT MSI from before and 10-months post-therapy (n=1).

Following these extensive functional studies, we treated the patient with low-dose tofacitinib (5mg daily). Remarkably, within eight weeks, circulating inflammatory markers (ESR and C-reactive protein) normalized (Figure 6D-E). Significant improvement in dermatitis followed, both grossly and histologically (Figure 6F). By six months, the patient reported complete resolution of gastrointestinal symptoms (decrease in modified PUCAI from 35-50 initially to 0). Biopsy of colonic tissue revealed restoration of crypt architecture and complete resolution of eosinophilic infiltrates (Figure 6G). Twelve months after the initiation of treatment, the patient remains stable and is awaiting re-transplantation.

Finally, we confirmed the pharmacological rescue of JAK hyperactivity in the patient’s cells after tofacitinib treatment. RNA isolated from PBMCs revealed that the expression of ISGs, which was significantly elevated before treatment, progressively declined to normal levels (Figure 6H). Mass cytometry analysis was then performed to confirm the decrease in basal STAT phosphorylation. Drastic reductions were observed across cell types in STAT3, STAT4, STAT5 and STAT6 phosphorylation, whereas STAT1 phosphorylation levels were reduced in some, but not all cell types (Figure 6I). Overall, these results validate S703I *JAK1* as the etiology *in vivo* of the widespread immune dysregulation, and they illustrate the striking power of precision medicine both as a life-saving approach for patients with rare diseases and as a means of discovering novel and fundamental physiological mechanisms.

## Discussion

Undiagnosed disease programs have proven remarkably successful at detecting potential causative variants of disease (Splinter *et al*., 2018; Lee *et al*., 2019; Yang *et al*., 2019). This report demonstrates the value of in-depth study of select patients identified in these programs. Most immediately, the findings described herein directed the successful treatment of a complex immunodysregulatory disease in a highly personalized molecular fashion. More broadly, this case implicates JAK1 dysfunction in common forms of multifactorial diseases, including dermatitis, enteritis, colitis and eosinophilic disorders. These features align with the other reported JAK1 GoF mutation recently published (Del Bel *et al*., 2017), and together, provide strong justification for the expanding use of JAK inhibitors in these disorders (O’Shea *et al*., 2015; O’Shea and Gadina, 2019). Yet, to date, MN has not been recognized to involve JAK-STAT dysregulation. It logically follows that MN, the most common cause of nephrotic syndrome, may be amenable to early treatment with JAK inhibitors. In fact, baricitinib, a JAK1/JAK2 inhibitor, demonstrated recent success in a clinical trial for diabetic nephropathy, a related nephrotic syndrome (Tuttle *et al*., 2018).

The absence of MN in the other reported JAK1 GoF mutation (A634D) may represent important distinctions in the behavior of different mutated forms of JAK1, rather than variable penetrance of the same genetic etiology. Disruptions of the pseudokinase domain may have vastly divergent functional consequences, which can already be gleaned by comparing the JAK1 mutations identified to date: P733L and P832S (LoF) vs. A634D and S703I (GoF). Moreover, our findings indicate that GoF may not necessarily lead to the universal activation of downstream pathways, as S703I caused the hyperactivation of some pathways, but not others. Furthermore, disruption of the pseudokinase domain by S703I enabled cells to respond promiscuously via non-canonical signaling pathways. Together, these findings suggest that the pseudokinase domain is not a simple “on/off” switch. This regulatory complexity may stem from the ability of the JAK1 pseudokinase to modulate the activity of JAK2 and TYK2, as demonstrated here. Consequently, careful study of each mutation is warranted, each yielding valuable information on fundamental JAK1 function. The complex regulation becomes especially important given the high incidence of oncogenic JAK mutations, as well as the expanding therapeutic use of JAK inhibitors. Regarding the latter, the evidence presented here and elsewhere (Haan *et al*., 2011; Li *et al*., 2013; Eletto *et al*., 2016) of the highly cooperative action of JAKs challenges the strategic wisdom of increasing the selectivity of JAK inhibitors.

Downstream from the JAK-receptor complex, S703I JAK1 upregulates STAT1, STAT2, STAT3, and STAT5 phosphorylation and, correspondingly, downstream gene transcription. We demonstrate here that this GoF occurs in a hyper-responsive fashion *ex vivo*, but that intrinsic signaling can occur independent of cytokine, as evidenced by basal STAT phosphorylation in the absence of a detectable increase in JAK-STAT cytokines. However, it is difficult to isolate which specific pathways cause disease, as JAK1 mediates signaling for more than twenty five cytokines (O’Shea *et al*., 2015). This broad immunodysregulatory disease is more likely to be driven by constitutive activity of a gamut of cytokine pathways, which have innumerable functions in autoinflammation and autoimmunity.

Nevertheless, specific immune disturbances were evident. We observed a robust upregulation of type I IFN signaling in this patient. Furthermore, in both published cases of JAK1 GoF eosinophils increase in circulation and infiltrate affected tissues. These findings are consistent with the features of atopic disease in both cases. Our high-dimensional analyses also detected a profound dysregulation of NK cells. CD56^hi^ NK cells demonstrated the largest difference in frequency, exhibited the most upregulated pSTAT profile, and carried the highest percentage of mutant transcript. Consistent with their phenotype, CD56^hi^ NK cells generally express high levels of *JAK1*, which likely potentiates the GoF effect in NK cells from the patient. This expansion likely arises from the intrinsic STAT5 activity and IL-2-hyperresponsive signaling of S703I JAK1, which is known to expand CD56hi NK cells (Poli *et al*., 2009). It is worth noting that long-term clinical treatment with IL-2 (Caligiuri *et al*., 1993) or IFNβ (Saraste, Irjala and Airas, 2007), disturbances that phenocopy JAK1 GoF, also lead to CD56^hi^ expansions. Interestingly, although tofacitinib clearly reduced transcriptomic and pSTAT inflammatory signatures and resolved clinical disease, we did not observe reversal of the CD56hi NK cell enrichment. The pharmacokinetic profile of tofacitinib may be such that it decreases the function of these cells, but not their number. In any case, it is apparent that persistent JAK1-mediated cytokine signaling has critical consequences on NK cells.

Finally, this study is among the first to demonstrate somatic mosaicism underlying immunologic disease (Holzelova *et al*., 2004; Saito *et al*., 2005; Wolach *et al*., 2005; Tanaka *et al*., 2011; Liu *et al*., 2014; de Inocencio *et al*., 2015). Mosaicism remains an underappreciated feature of monogenic disorders. Recent studies indicate that undetected somatic variation is incredibly common (predicted 10^16^ base substitutions), and that up to 6.5% of *de novo* mutations identified to date are actually mosaic (Acuna-Hidalgo *et al*., 2015). Mosaicism becomes especially important as we incorporate WES into the future of clinical medicine. Potential disease-causing mutations may be overlooked without careful consideration of the tissue site genotyped, loci with low frequency variants, and the poor sensitivity of classic Sanger sequencing. Additionally, asymmetric clinical manifestations like those observed here (leg length discrepancy and dermatitis along lines of embryological migration) should prompt suspicions of mosaicism and guide genetic analysis.

The analyses described here have only recently become possible with technological advances in single-cell assays. Here, by adapting the inDrop single cell RNA-seq platform, we implement a novel approach for the detection and analysis of a specific mutation within a transcript region not readily accessible by other methods. The resulting single cell resolution data allowed us to determine mutant allele frequency in different cell populations, identify mutation-associated gene expression patterns, and unexpectedly, to observe allelic bias in transcription of *JAK1*. This bias is consistent with recent evidence of widespread transcriptional bursting and monoallelic expression of autosomal genes (Gimelbrant *et al*., 2007; Jeffries *et al*., 2012; Borel *et al*., 2015; Reinius and Sandberg, 2015). Importantly, this is the first report of monoallelic expression of a mutated gene. Biased allelic expression, in conjunction with mosaicism, may prove an important point of focus for future genetic studies of variable penetrance, affected carriers and undiagnosed disease.

## Acknowledgements

We thank Adeeb Rahman, Brian Lee, Kevin Tuballes and Laura Walker from Human Immune Monitoring Center and Rachel Brody from the Biorepository and Pathology CoRE at the Icahn School of Medicine for their technical assistance and guidance in experimental design. This research was supported by National Institute of Allergy and Infectious Diseases Grants R01AI127372, R21 AI134366 and R21AI129827, and funding from the March of Dimes, awarded to DB. Research in the Mace laboratory was supported by JSO R01AI120989 and research in the Rosenberg laboratory by DP5 OD012142. CG was supported by T32 training grant 5T32HD075735-07.

## Author contributions

CG designed and performed most of the experiments, analyzed the data and wrote the manuscript. JC analyzed all scRNAseq data and edited the manuscript. SB generated cell lines and processed whole blood experiments. GE analyzed sequencing data, performed ddPCR and edited the manuscript. JM performed the tissue processing for ddPCR. SU and RC designed the platform and carried out the custom scRNAseq experiments. EM and JO performed the NK cell experiments and analysis. BW, MM, BG helped to design the experiments and analyze data. LJ, DD, RP, JS performed clinical analyses and reports. BR designed the custom scRNAseq, analyzed the data and edited the manuscript. BW, MM, JO, BG helped to design the experiments and analyze data. DB helped to design the experiments and analyze data, supervised the work and wrote the manuscript. All authors commented on the manuscript.

## Data Availability

The raw data for experiments performed will be made available, including the PCR, multiplex ELISA, mass cytometry, flow cytometry, scRNAseq data shown in Figure 1D, 2C, 4, 5, 6C-I, S1E, S3, S4A, S4C, S5A-C, S6, S7. Web links will be made available for large files, including fcs files and scRNAseq gene x cell matrices. For privacy concerns of the study participants, the complete data files from whole exome sequencing will be restricted to the variants in Table 1.

## Code Availability

Python scripts and associated Seurat code (R-based) used for the tailored analysis of JAK1-specific scRNAseq will be made available upon publication of the manuscript. Contact.

