## Supplemental Files and Text for "Complex Autoinflammatory Syndrome Unveils Fundamental Principles of *JAK1* Transcriptional and Biochemical Function"

### Supplemental Files for *Complex Autoinflammatory Syndrome Unveils Fundamental Principles of JAK1 Transcriptional and Biochemical Function*

#### Contents


1. Supplementary Case Report
2. Supplemental Figures and Tables
3. Figure Legends for Supplemental Figures
4. Methods
5. Supplemental References

**SUPPLEMENTARY CASE REPORT**

18 year-old female who presented during early childhood with persistent, recurrent cutaneous and gastrointestinal inflammatory disease with eosinophilic infiltration and peripheral eosinophilia. Notable physical features include a leg length discrepancy, short stature and low body weight. In parallel, she developed refractory membranous nephropathy leading to end stage renal disease. A kidney transplant was complicated by disease recurrence in the graft, progressing over several years, as well as acute rejection ultimately rendering the patient dialysis-dependent. Specifically, her past medical history by organ involvement is further details as follows:

**Renal**: At 3 years of age, she developed rapid weight gain, edema and proteinuria. Renal biopsy at age 7 demonstrated membranous nephropathy, which was refractory to treatment with corticosteroids, and later cyclosporine and tacrolimus. By history the nephrotic syndrome was ameliorated by use of an elemental diet but this was not able to be consistently maintained. Serological testing for anti-PLA-2R receptor, anti-thrombospondin and anti–bovine serum albumin were all negative. A gradual decline in renal function was observed and at age 11 a living-donor kidney transplantation was performed. One year later, nephrotic range proteinuria recurred, and biopsy confirmed relapsing membranous nephropathy. The graft function further declined, and an episode of acute antibody-mediated rejection resulted in transplant failure at age 16. She has required long-term hemodialysis since that time, delivered via an AV fistula, and is currently being evaluated for a second transplant.

**Dermatologic**: At birth, a pustular rash involving face and extremities was noted. The rash persisted and worsened after discharge. Skin biopsy at 3 months suggested Inflammatory Linear Verrucous Nevus. Skin involvement spread and worsened in intensity with age, manifesting as a diffuse, erythematous rash involving the face, trunk and extremities, with prominence on the left side. Biopsies later demonstrated a subacute or chronic spongiotic dermatitis. The epidermis was acanthotic and showed varying degrees of intercellular edema (spongiosis) with widening of the intercellular spaces; the stratum corneum was thickened and focally compact; the dermis contained a perivascular lymphohistiocytic infiltrate which extended around the superficial and deep vascular plexus. Of note, the biopsies did not show the changes commonly associated with epidermal nevi: alternating ortho and parakeratosis, epidermolytic hyperkeratosis, or acantholytic dyskeratosis. Rather, these clinical and histologic changes in the skin likely represent a form of blaschkitis, an inflammatory skin condition, presenting as papules or vesicles, occurring along the lines of Blaschko (which represent somatically distinct bands of ectodermal migration).

**Gastrointestinal**: In infancy, she experienced recurrent emesis and diarrhea unresponsive to formula changes. Bloody stools were noted at 10 months of age and watery diarrhea and abdominal pain became persistent. Repeat endoscopic biopsies demonstrated chronic, unspecified inflammation at various sites (most frequently colonic, but also gastric, duodenal, ileal and esophageal regions). Eosinophilic infiltration of the colon was consistently noted. Symptoms were only marginally responsive to treatment with corticosteroids, chronic antibiotics and a severely restricted diet.

**Growth disturbances**: Growth impairment was reflected by short stature (Z score <-3) and low body weight (Z score -2 to -8), currently 138 cm and 31 kg. Nutritional etiologies were addressed by placement of a G-tube at age 10 with some improvement in growth. Growth hormone was administered for 5 years with moderate benefit. Leg length discrepancy was identified at birth, with left extremity smaller than right in girth and length.

**Immunologic:** Allergic reactions were observed to enalapril (anaphylaxis), milk (rash), soy (rash) and wheat (rash). She also experienced occasional episodes of dyspnea along with lip and leg angioedema, without an identifiable inciting allergen. Asthma was diagnosed and managed with bronchodilators. Acute phase reactants were noted to be consistently elevated, including ESR and C-reactive protein. Complement (C3 and C4) levels were within reference range. Likewise, quantitative immunoglobulin testing for IgG, IgA, IgE and IgM was within normal limits. Seroconversion after immunization was observed for all vaccinations except varicella virus, hepatitis A virus and hepatitis B virus.

Family History was largely unremarkable, with no family history of consanguinity or gastrointestinal, renal, immunologic or dermatologic disease. Mother, father and older brother are alive and well.

**Response to tofacitinib**
After 8 weeks of tofacitinib there was complete normalization of acute phase reactants, ESR and C-reactive protein upon laboratory assessment. The patient was previously unable to tolerate dairy, soy and gluten due to severe abdominal pain and diarrhea. After initiation of tofacitinib, she liberalized her diet without restriction and remained asymptomatic, with formed stools and without abdominal pain.

At baseline the endoscopic findings included altered vascularity and friable mucosa from rectum to descending colon with microscopic patchy, active colitis with eosinophilic infiltration noted in the ascending colon. After 6 months of treatment on tofacitinib 5mg daily, the colon was grossly normal, with microscopic active colitis and complete resolution of eosinophilia. The dose was further increased to 7.5 mg daily thereafter. The dermatitis significantly improved, as seen in the images.

Tofacitinib is 30% renally-excreted and the dose administered was limited due to chronic kidney disease. Presumably, after re-transplantation the dose may be escalated with a potentially greater effect. To date, the drug has been very well tolerated with no evidence of adverse effects with close monitoring.

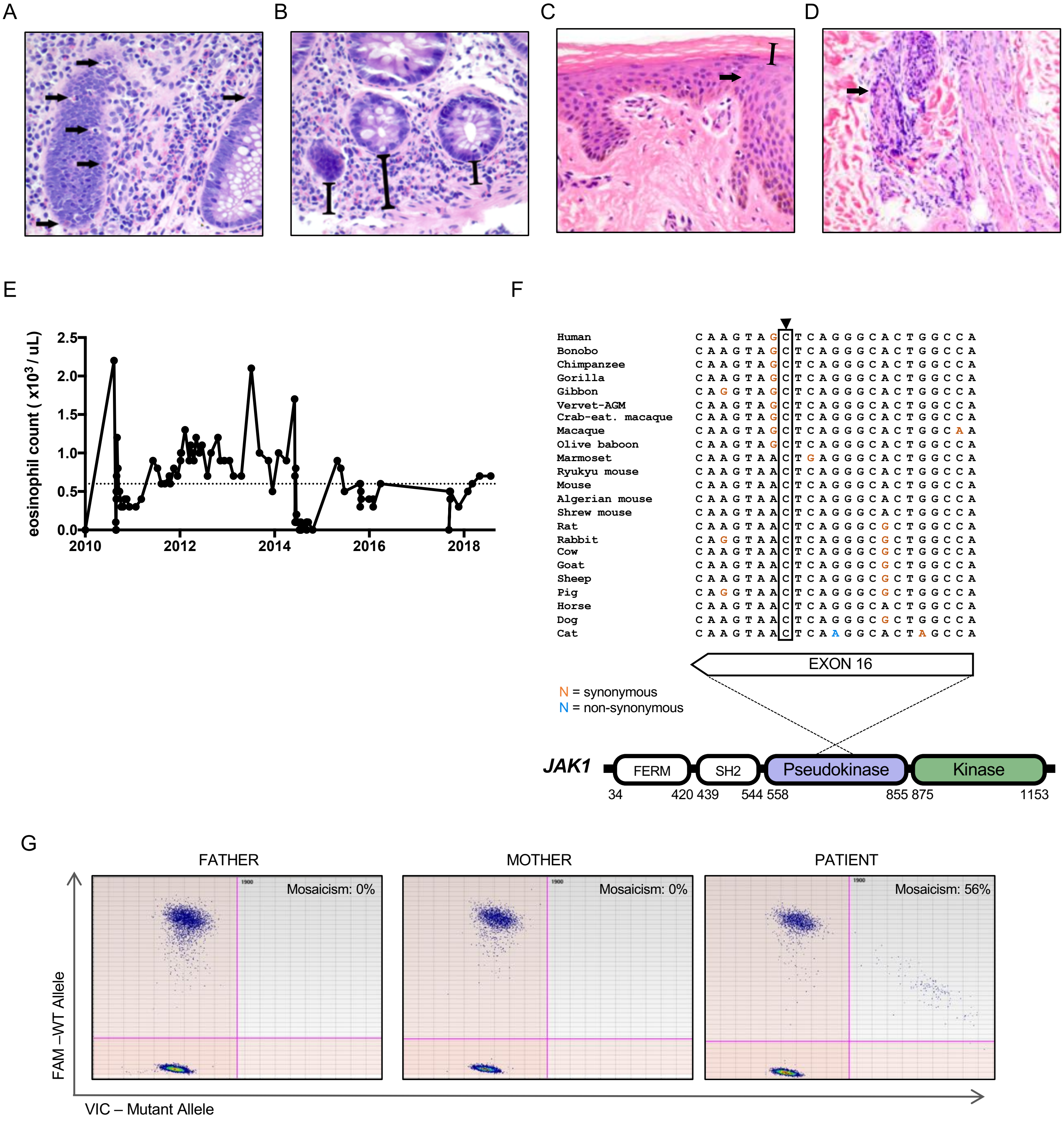


**Supplementary Figure 1. Whole exome sequencing uncovers a novel JAK1 mutation in a patient with complex immune dysregulation.**

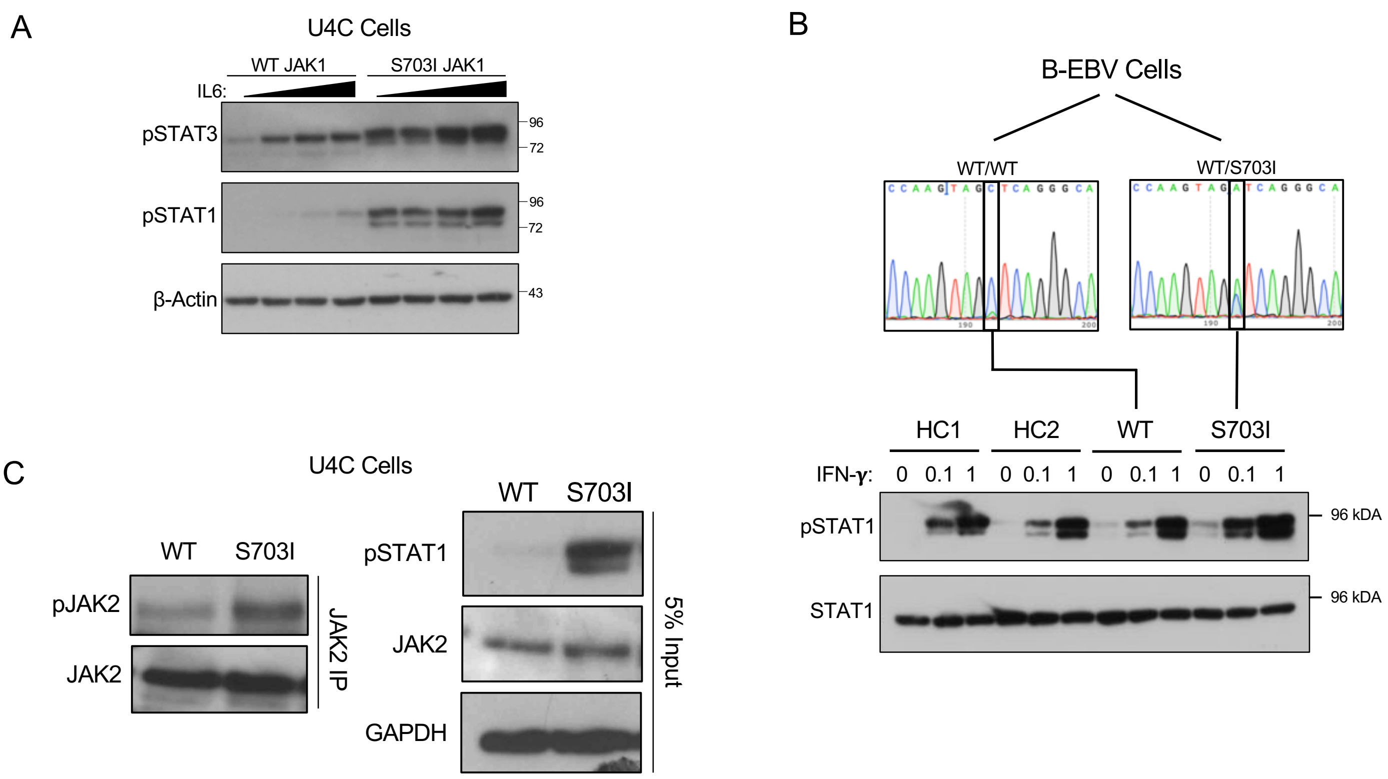


**Supplementary Figure 2.** ***In vitro*** **analysis of S703I JAK1 demonstrates basal and hyper-active JAK-STAT signaling.**

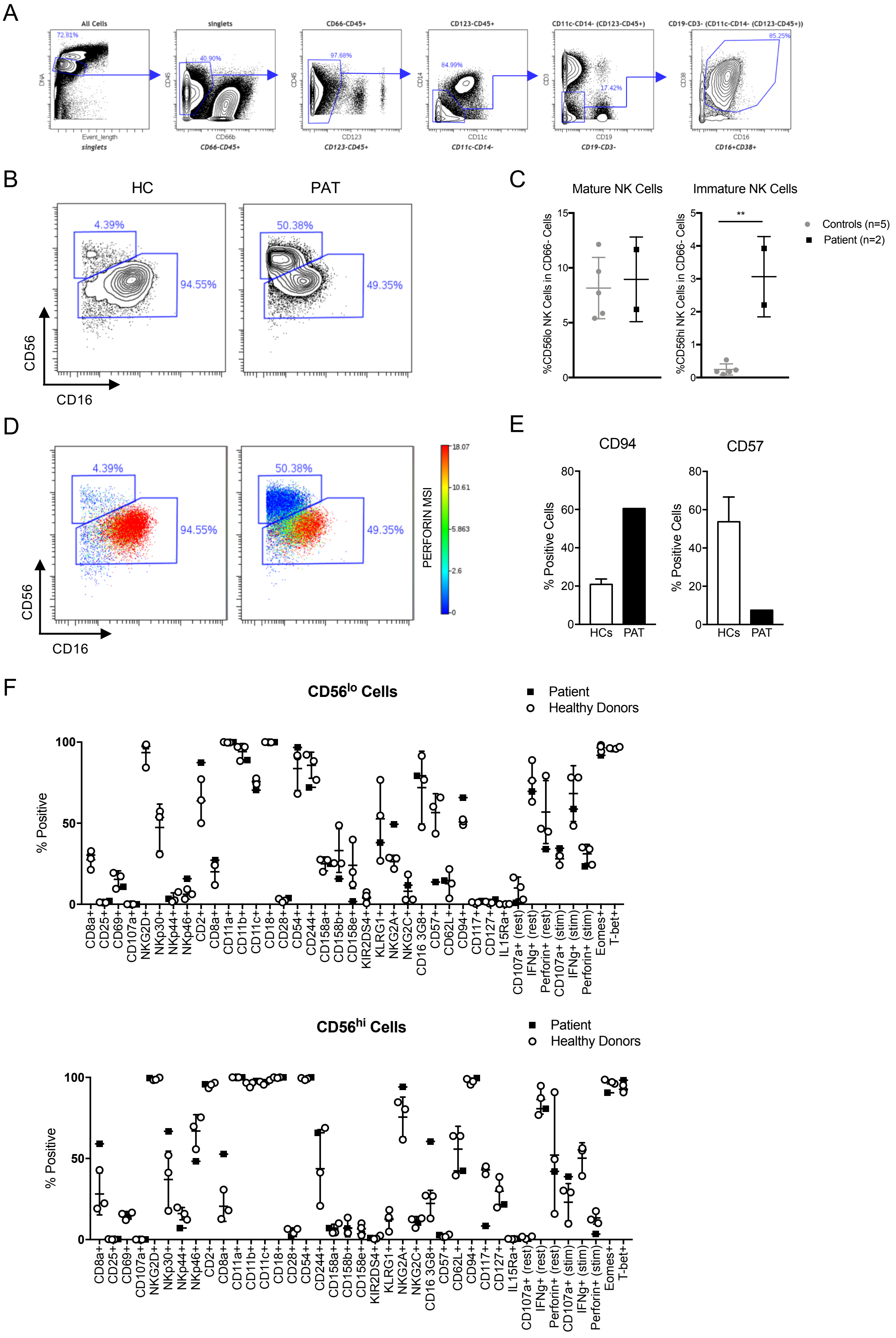


**Supplementary Figure 3. Detailed immunophenotyping of patient NK cells.**

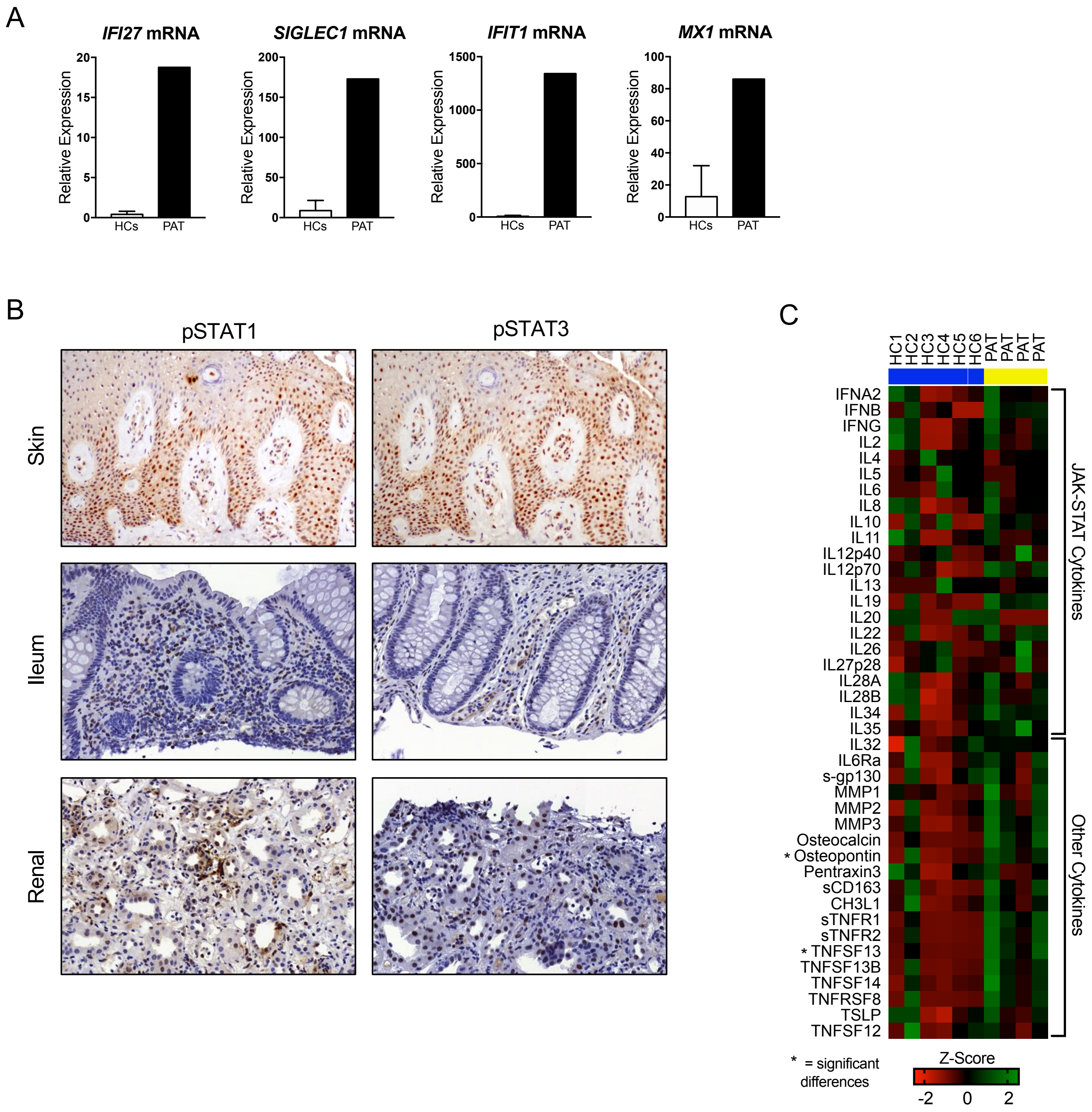


**Supplementary Figure 4. Characterization of *in vivo* JAK-STAT signaling.**

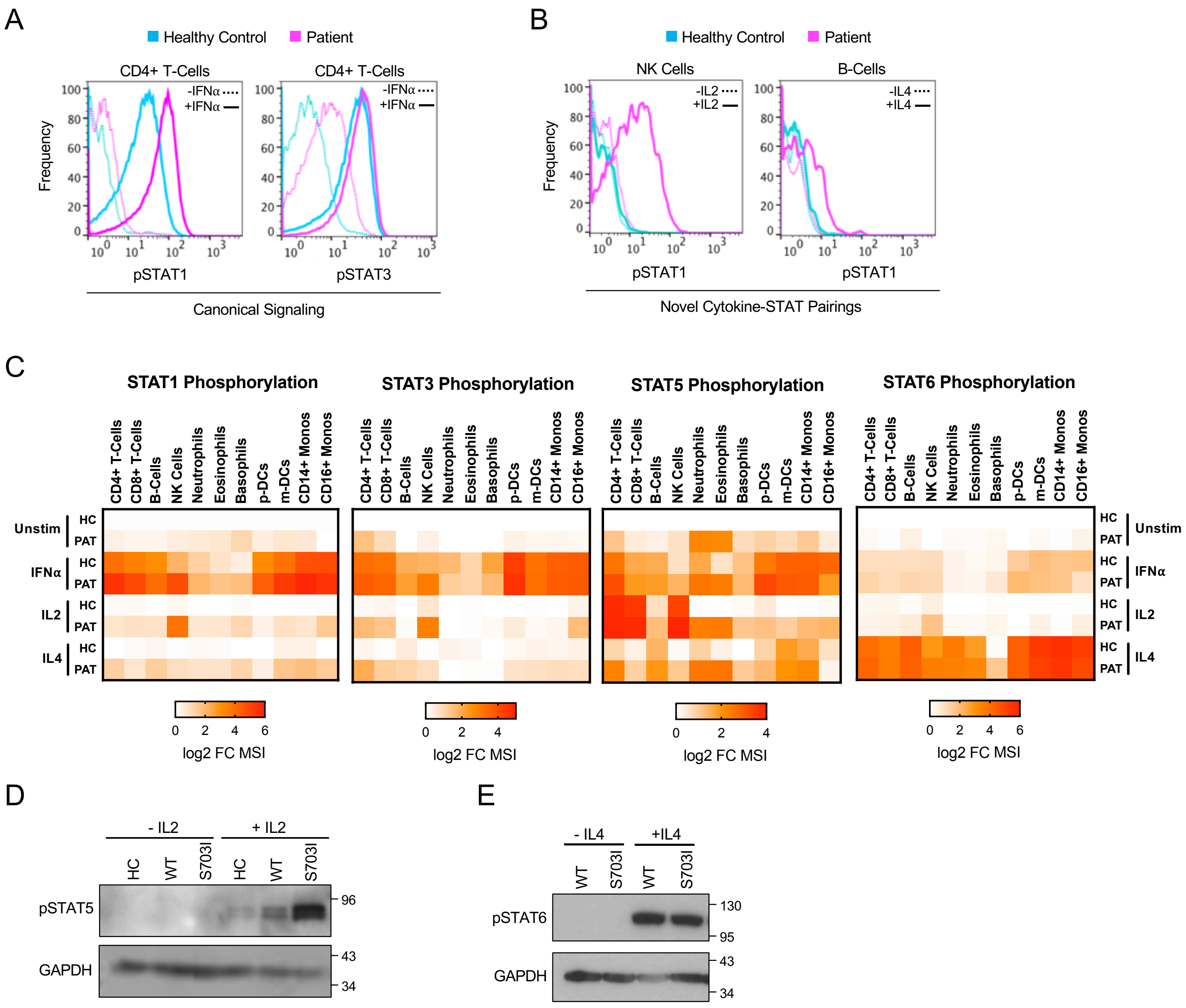


**Supplementary Figure 5. STAT phosphorylation responses to cytokine stimulation.**

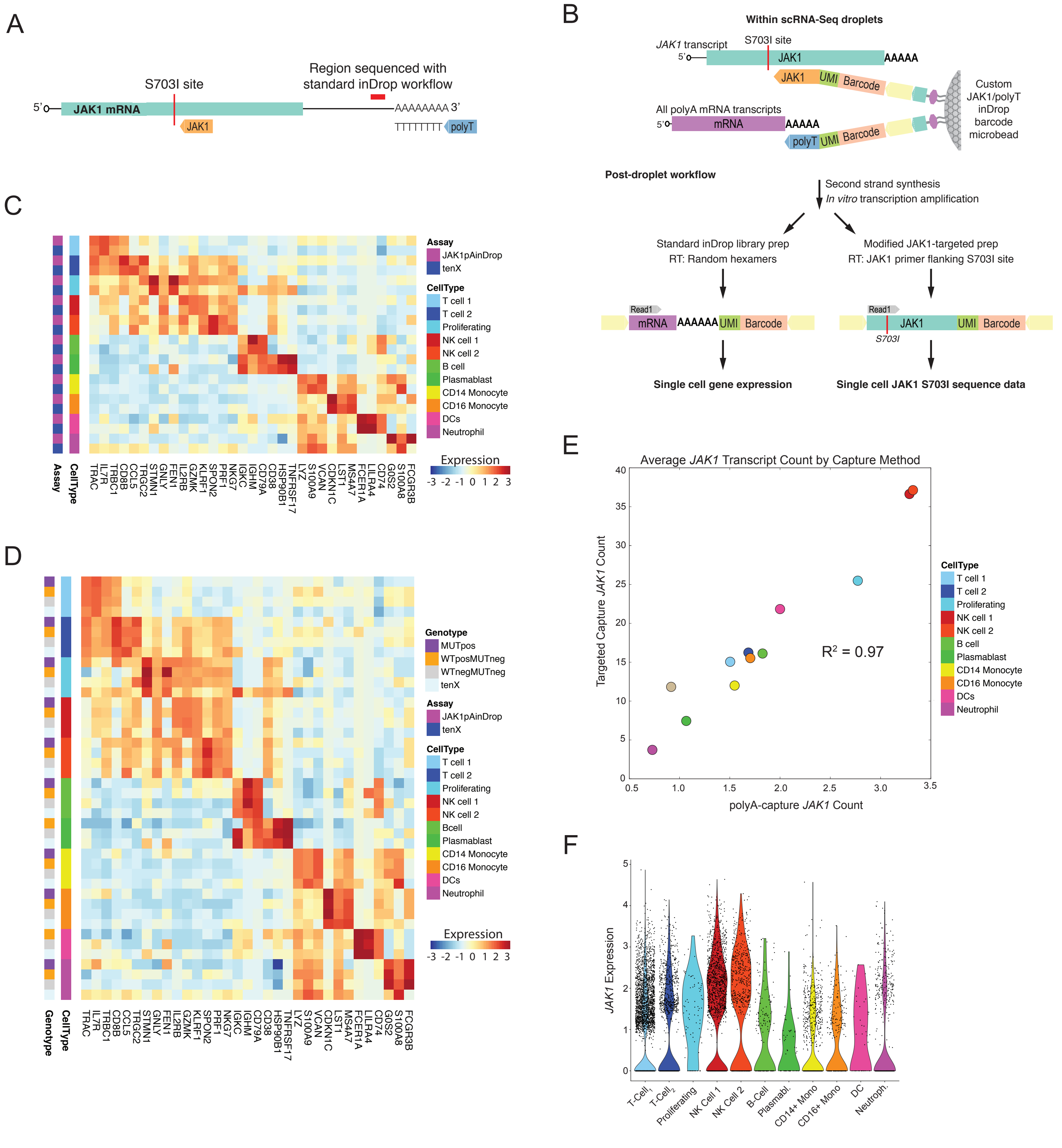


**Supplementary Figure 6. Adapted scRNAseq platform maps genetic distribution and transcriptomic signatures of S703I JAK1.**

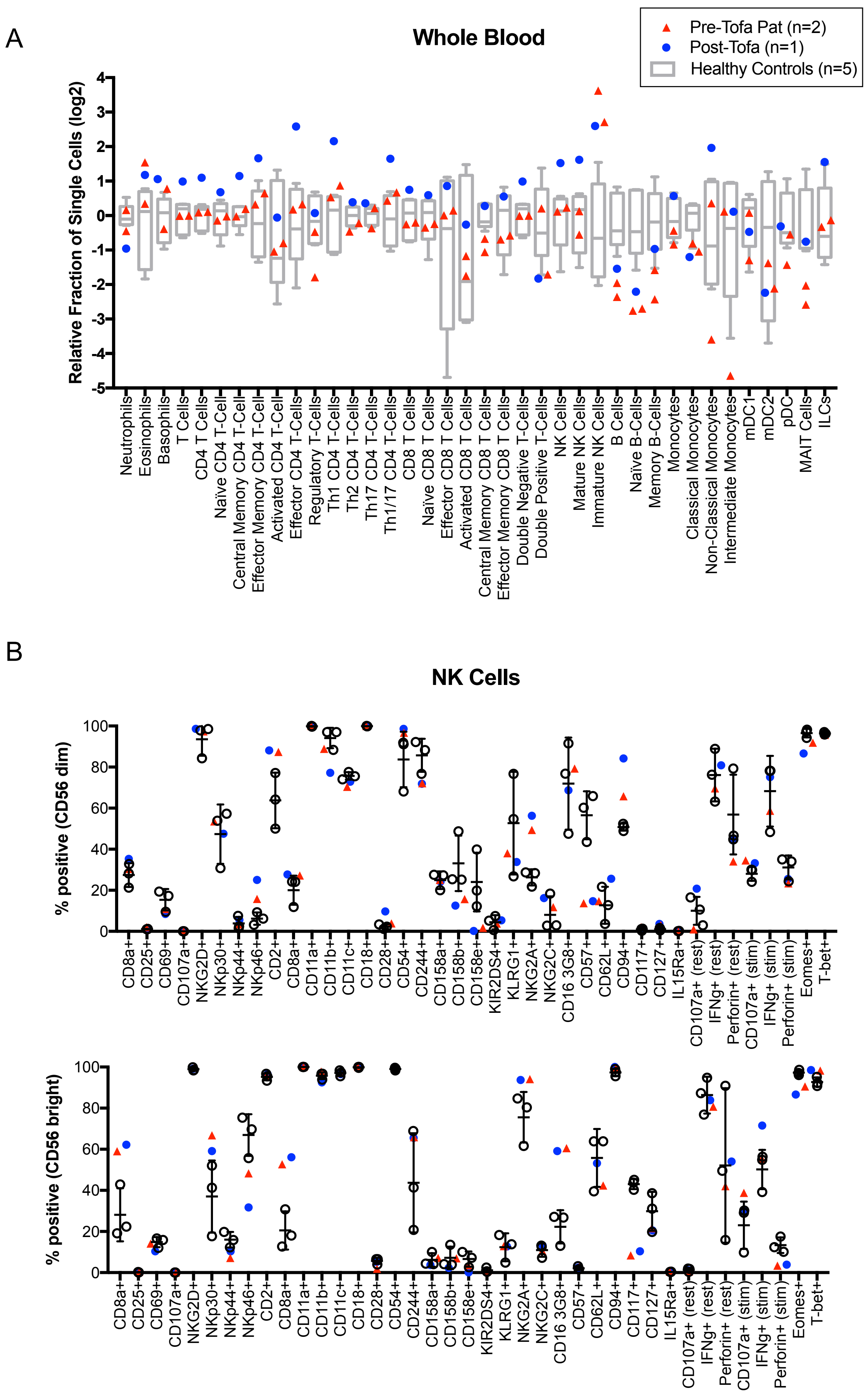


**Supplementary Figure 7. CyTOF and NK Cell immunophenotyping before and after tofacitinib therapy.**

| Recessive Inheritance Pattern | | | | | | | | | |
|---|---|---|---|---|---|---|---|---|---|
| Gene Symbol | Patient Allele Fraction | Maternal Allele Fraction | Maternal Allele Fraction | Protein Variant | Translation Impact | CADD | MSC Threshold | PolyPhen-2 Function Prediction | Inferred Activity |
| *ANKRD36C* | 15.48 | 21.62 | 14.81 | p.Q642fs / p.Q75fs | frameshift | <10 | 3.3 | N/A | Loss-of-function |
| *LYZ* | 8.18 | 12.59 | 16 | p.A65fs / p.G66fs | frameshift | 20.2 | 23.7 | N/A | Loss-of-function |

| *De Novo* Inheritance Pattern | | | | | | | | | |
|---|---|---|---|---|---|---|---|---|---|
| Gene Symbol | Patient Allele Fraction | Maternal Allele Fraction | Maternal Allele Fraction | Protein Variant | Translation Impact | CADD | MSC Threshold | PolyPhen-2 Function Prediction | Inferred Activity |
| *JAK1* | 0.28 | 0.00 | 0.00 | p.S703I | missense | 27.6 | 3.3 | Probably Damaging | Gain of Function |
| *IFNA14* | 0.12 | 0.00 | 0.00 | p.K144R | missense | <10 | 3.3 | Benign | Unknown |

| Dominant Inheritance Pattern | | | | | | | | | |
|---|---|---|---|---|---|---|---|---|---|
| Gene Symbol | Patient Allele Fraction | Maternal Allele Fraction | Maternal Allele Fraction | Protein Variant | Translation Impact | CADD | MSC Threshold | PolyPhen-2 Function Prediction | Inferred Activity |
| *MTOR* | 0.50 | 0.00 | 0.52 | p.Y1913C | missense | 28.8 | 24.6 | Probably Damaging | Haplo-insufficiency |
| *GFPT1* | 0.47 | 0.00 | 0.37 | p.R163W | missense | 33 | 0.0 | Possibly Damaging | Normal |
| *DYSF* | 0.39 | 0.00 | 0.48 | p.R762C | missense | 35 | 0.0 | Probably Damaging | Normal |
| *LRP2* | 0.54 | 0.56 | 0.00 | p.Y1681H | missense | <10 | 4.8 | Possibly Damaging | Unknown |
| *ALPP* | 0.43 | 0.00 | 0.43 | p.G148R | missense | 24.6 | 3.3 | Probably Damaging | Normal |
| *RHOA* | 0.14 | 0.15 | 0.00 | p.A44A | splice site loss | <10 | NA | - | Normal |
| *HLA-B* | 1.00 | 1.00 | 0.00 | p.V127M | promoter loss; missense | <10 | 3.3 | Benign | Unknown |
| *NOM1* | 0.54 | 0.54 | 0.00 | p.R782Q | missense | 32 | 3.3 | Probably Damaging | Haplo-insufficiency |
| *PLEC* | 0.51 | 0.00 | 0.50 | p.T2912S | missense | 23.3 | 7.2 | Possibly Damaging | Normal |
| *WNK1* | 0.50 | 0.39 | 0.00 | p.I801V | missense | <10 | 0.0 | Benign | Unknown |
| *COL4A1* | 0.54 | 0.00 | 0.53 | p.P784L | missense | 13.6 | 0.2 | Probably Damaging | Haplo-insufficiency |
| *MEFV* | 0.44 | 0.00 | 0.60 | p.A744D | missense | 10.83 | 0.0 | Benign | Unknown |
| *SNTB2* | 0.36 | 0.00 | 0.55 | p.L262M | missense | 24.9 | 9.3 | Possibly Damaging | Unknown |
| *SOCS3* | 0.49 | 0.00 | 0.45 | p.T208S | missense | 13.46 | 5.4 | Benign | Unknown |
| *RYR1* | 0.55 | 0.51 | 0.00 | p.Y1121C | missense | 27 | 0.2 | Probably Damaging | Haplo-insufficiency |
| *PRODH* | 0.52 | 0.46 | 0.00 | p.V554M | missense | 34 | 7.6 | Probably Damaging | Haplo-insufficiency |
| *POLA1* | 0.47 | 0.43 | 0.00 | p.P496S | missense | 27.2 | 3.3 | Possibly Damaging | Haplo-insufficiency |
| *IKBKG* | 0.52 | 0.00 | 1.00 | p.E128D | missense | 15.28 | 0.0 | Benign | Normal |

**Table 1. Variants of interest identified from whole exome sequence and variant analysis by models for recessive, *de novo* and dominant inheritance.**

| Cell Type | CyTOF Markers | Healthy Control | | Patient | |
|---|---|---|---|---|---|
| **Granulocytes** | | | | | |
| Neutrophils | CD45+CD66b+CD16hi | 40.17 - 70.40 | % | 25.16 - 54.62 | % |
| Eosinophils | CD45+CD66b+CD16hi | 0.90 - 5.52 | % | 4.08 - 9.38 | % |
| Basophils | CD45+CD66b-123+CD11c-CD14-CD3-CD20-CD38+HLADR- | 0.25 - 0.78 | % | 0.38 - 1.02 | % |
| **B Cells** | | | | | |
| Total | CD45+CD66b-CD123-CD11c-CD14-CD3-CD19+ | 1.29 - 5.20 | % | 0.57 - 1.01 | % |
| Naiv | CD45+CD66b-CD123-CD11c-CD14-CD3-CD19+CD27-IgD+ | 0.66 - 3.32 | % | 0.29 - 0.43 | % |
| Memory | CD45+CD66b-CD123-CD11c-CD14-CD3-CD19+CD27+ | 0.19 - 1.19 | % | 0.10 - 0.28 | % |
| **Dendritic Cells** | | | | | |
| mDC | CD45+CD66b-CD123-CD11c-CD14-CD3-CD19-HLADR+CD14-CD11c+CD11b+ | 0.02 - 0.12 | % | 0.03 - 0.07 | % |
| mDC1 | CD45+CD66b-CD123-CD11c-CD14-CD3-CD19-HLADR+CD14-CD11c+CD11b+CD1c+ | 0.02 - 0.09 | % | 0.03 - 0.06 | % |
| mDC2 | CD45+CD66b-CD123-CD11c-CD14-CD3-CD19-HLADR+CD14-CD11c++CD11b+CD1c+CD141- | 0.00 - 0.03 | % | 0.00 - 0.00 | % |
| pDC | CD45+CD66b-123+CD11c-CD14-CD3-CD20-CD38+HLADR+ | 0.08 - 0.32 | % | 0.06 - 0.12 | % |
| **Monocytes** | | | | | |
| Total | CD45+CD66b-CD123-CD3-CD19-HLADR+CD14+CD11c+CD14+ | 2.77 - 10.37 | % | 2.12 - 9.70 | % |
| Classical | CD45+CD66b-CD123-CD3-CD19-HLADR+CD14+CD11c+CD14hiCD16- | 2.33 - 5.69 | % | 1.95 - 2.56 | % |
| Non-Classical | CD45+CD66b-CD123-CD3-CD19-HLADR+CD14loCD16hiCD11c+CX3CR1 | 0.38 - 3.41 | % | 0.14 - 6.43 | % |
| Intermediate | CD45+CD66b-CD123-CD3-CD19-HLADR+CD14+CD11c+CD14hiCD16+ | 0.06 - 1.27 | % | 0.03 - 0.71 | % |
| **NK Cells** | | | | | |
| Total | CD45+CD66b-CD123-CD11c-CD14-CD3-CD19-CD38+CD16+ | 1.37 - 6.28 | % | 3.32 - 13.19 | % |
| Mature | CD45+CD66b-CD123-CD11c-CD14-CD3-CD19-CD38+CD56loCD16+ | 1.35 - 5.93 | % | 2.61 - 11.74 | % |
| Immature | CD45+CD66b-CD123-CD11c-CD14-CD3-CD19-CD38+CD56loCD16lo | 0.03 - 0.34 | % | 0.71 - 1.45 | % |
| **CD4 T Cells** | | | | | |
| Total | CD45+CD66b-CD123-CD11c-CD14-CD19-CD3+CD4+ | 9.02 - 16.14 | % | 13.78 - 27.72 | % |
| Naïve | CD45+CD66b-CD123-CD11c-CD14-CD19-CD3+CD4+CD27+CD45RA+ | 2.77 - 6.96 | % | 4.63 - 8.18 | % |
| Effector | CD45+CD66b-CD123-CD11c-CD14-CD19-CD3+CD4+CD27-CD45RA+ | 0.03 - 0.41 | % | 0.08 - 0.16 | % |
| Activated | CD45+CD66b-CD123-CD11c-CD14-CD19-CD3+CD4+CD38+HLADR+CD25+ | 0.01 - 0.07 | % | 0.04 - 0.22 | % |
| Central Memory | CD45+CD66b-CD123-CD11c-CD14-CD19-CD3+CD4+CD45RA-CD38-HLADR-CD27+CD197+ | 4.05 - 7.13 | % | 5.86 - 13.15 | % |
| Effector Memory | CD45+CD66b-CD123-CD11c-CD14-CD19-CD3+CD4+CD45RA-CD38-HLADR-CD27+CD197- | 0.39 - 1.99 | % | 1.23 - 3.11 | % |
| Th1 | CD45+CD66b-CD123-CD11c-CD14-CD19-CD3+CD4+CCR6-CXCR3+CCR4- | 1.02 - 3.37 | % | 3.24 - 10.01 | % |
| Th2 | CD45+CD66b-CD123-CD11c-CD14-CD19-CD3+CD4+CCR6-CXCR3-CCR4+ | 5.48 - 8.54 | % | 5.11 - 9.23 | % |
| Th17 | CD45+CD66b-CD123-CD11c-CD14-CD19-CD3+CD4+CCR6+CXCR3-CCR4+(CD161+) | 1.34 - 2.08 | % | 1.31 - 2.16 | % |
| Th1/17 | CD45+CD66b-CD123-CD11c-CD14-CD19-CD3+CD4+CCR6+CXCR3+CCR4- | 1.04 - 3.45 | % | 2.88 - 6.70 | % |
| CD8 T Cells | | | | | |
| Total | CD45+CD66b-CD123-CD11c-CD14-CD19-CD3+CD8+ | 4.57 - 10.75 | % | 6.51 - 13.02 | % |
| Naïve | CD45+CD66b-CD123-CD11c-CD14-CD19-CD3+CD8+HLADR-CD45RA+CD27+ | 1.30 - 4.33 | % | 2.48 - 4.77 | % |
| Effector | CD45+CD66b-CD123-CD11c-CD14-CD19-CD3+CD8+HLADR-CD45RA+CD27- | 0.05 - 2.91 | % | 1.35 - 2.44 | % |
| Activated | CD45+CD66b-CD123-CD11c-CD14-CD19-CD3+CD8+HLADR+ | 0.03 - 0.72 | % | 0.08 - 0.22 | % |
| Central Memory | CD45+CD66b-CD123-CD11c-CD14-CD19-CD45RA-CD38-HLADR-CD27+CD197+ | 0.87 - 1.53 | % | 0.57 - 1.45 | % |
| Effector Memory | CD45+CD66b-CD123-CD11c-CD14-CD19-CD45RA-CD38-HLADR-CD27+CD197- | 0.45 - 2.61 | % | 0.91 - 2.16 | % |
| Other | | | | | |
| MAIT Cells | CD45+CD66b-CD123-CD19-CD161+CD3+CD4-CD8any | 1.15 - 5.91 | % | 0.39 - 1.38 | % |
| ILCs | CD45+CD66b-CD123-CD3-CD19-CD14-CD11c-CD161+CD127+ | 0.01 - 0.04 | % | 0.01 - 0.04 | % |

**Table 2. Immune cell populations identified from whole blood by mass cytometry and range in healthy controls (n=5) and patient (2 replicates).**

**Supplementary Figure 1. Whole exome sequencing uncovers a novel JAK1 mutation in a patient with complex immune dysregulation.** (A) Histology of the cecal mucosa showing expansion of the lamina propria secondary to increased inflammatory cell infiltrate, with eosinophils in the lamina propria and crypt epithelium (arrows). (B) Colonic biopsy from right colon demonstrating accumulation of inflammatory cells beneath the crypts, a sign of chronicity ("crypt shortfall") (bars) (C) Skin biopsies demonstrating subacute or chronic spongiotic dermatitis with widening of the intercellular spaces (arrow) and thickening and focal compaction of the stratum corneum (bars). (D) Additional biopsy of the dermis showing perivascular lymphohistiocytic infiltrate extending around the superficial and deep vascular plexus (arrow). (E) Eosinophil count from peripheral blood. Dotted line represents upper limit of normal (F). Nucleotide alignment for exon 16 of *JAK1* across mammalian species, with synonymous (orange) and non-synonymous (blue) changes noted. (G) Digital droplet PCR of PBMC genomic DNA using mutation specific probes.

**Supplementary Figure 2. *In vitro* analysis of S703I JAK1 demonstrates basal and hyper-active JAK-STAT signaling.** (A) IL6 stimulation of transduced U4C cells at increasing concentrations (0, 5, 10, or 25 ng/mL). (B) Single cell cloning of patient-derived B-EBV cells into WT/WT and WT/S703I clones, and subsequent analysis of response to Type II IFN (ng/mL). (C) Immunoprecipitation of JAK2 and analysis of phosphorylation at the activation loop by immunoblotting from transduced U4C cells.

**Supplementary Figure 3. Detailed immunophenotyping of patient NK cells.** (A) Representative gating of NK cells from immunophenotyping by mass cytometry. (B) Analysis of NK cells by CD56 expression in patient and representative control. (C) Quantification of mature ($CD56^{lo}$) and immature ($CD56^{hi}$) NK cells as percent of CD66- peripheral blood cells from patient (two timepoints) and 5 healthy controls. (D) Median signal intensity (MSI) of perforin in NK cells from patient and a representative control. (E) Fraction of NK cells positive for CD94 and CD57 by flow cytometry of patient and control (n=3) PBMCs. (F) Percent positive for phenotypical and functional NK cell markers from $CD56^{lo}$ NK cells (top) and $CD56^{hi}$ NK cells (bottom) from patient and controls (n=3) PBMCs.

**Supplementary Figure 4. Characterization of *in vivo* JAK-STAT signaling.** (A) mRNA expression of interferon-stimulated genes from bulk PBMC of patient and healthy controls (n=3). (B) Immunohistochemistry from patient biopsies of indicated tissues stained with antibodies against phospho-STAT1 or phospho-STAT3 (brown) and counterstain (blue). (C) Multiplex ELISA for indicated cytokines from plasma of patient (4 replicates) and 6 healthy controls. Color intensity represents Z-score for cytokine concentrations.

**Supplementary Figure 5. STAT phosphorylation responses to cytokine stimulation.** (A) Representative histograms demonstrating basal (dotted lines) and ex vivo cytokine-stimulated (solid lines) phospho-STAT levels in CyTOF analysis of whole blood immune cells. (B) Histograms representing promiscuous phosphorylation of STAT1 following *ex vivo* IL-2 and IL-4 stimulation of whole blood. (C) Composite CyTOF data from Figure 4C-D represented as log2 fold change over the MSI of healthy control. (D) Immunoblotting for STAT5 phosphorylation after 15m stimulation with IL-2 (50 ng/mL) in healthy control, patient WT, and patient mutant B-EBV cells. (E) Immunoblotting for STAT6 phosphorylation in B-EBV cells stimulated with IL-4 (50 ng/mL) for 15m.

**Supplementary Figure 6. Adapted scRNAseq platform maps genetic distribution and transcriptomic signatures of S703I JAK1.** (A) Diagram illustrating the inability of standard inDrop platforms to capture the S703I site on *JAK1* mRNA. (B) Schematic of custom inDrop scRNAseq methodology adapted to target exon 16 of JAK1 platform, diagramming the capture platform and post-droplet workflow. (C) Heatmap of cell types defined in *JAK1*-specific inDrop platform and control 10X Chromium scRNAseq, demonstrating concordance of cell types. (D) Cell type marker expression across cells genotyped by identity of *JAK1* transcript detected. Cells containing <5 unique transcripts (gray), cells containing exclusively WT *JAK1* transcript (orange), or cells containing >5 S703I *JAK1* transcript. (E) Correlation of average count of unique *JAK1* transcripts captured by standard poly-T primers as compared to enriched capture by JAK1-exon-16-specific primers. (F) Total *JAK1* expression from poly-T capture across different cell types. Expression represented as log((TPM+100)/100).

**Supplementary Figure 7. CyTOF and NK Cell immunophenotyping before and after tofacitinib therapy. (A)** Frequency of whole blood immune cell subsets relative to healthy controls (gray, n=5) in patient cells, before (red) or after 3 months treatment with tofacitinib (blue). (B) Flow cytometry of $CD56^{lo}$ and $CD56^{hi}$ NK cells for phenotypical and functional NK cell markers. Healthy controls (open circles) n=3. Patient cells before (red) or after 3 months treatment with tofacitinib (blue).

**Table 1. Whole Exome Sequencing Variant Analysis**. Variants of interest identified from whole exome sequence and variant analysis by models for recessive, *de novo* and dominant inheritance.

**Table 2. Mass Cytometry Immunophenotyping of Peripheral Blood**. Immune cell populations identified from whole blood by mass cytometry and the quantification of the frequency range in healthy controls (n=5) and patient (2 replicates) as expressed by % of single cells.

#### Methods

##### Variant analysis

All samples were collected with informed consent in accordance with IRB-approved protocols (Study ID# IF2349568). For whole-exome sequencing, DNA was isolated (Qiagen Cat No 69504) from Ficoll-isolated granulocytes from whole blood of the proband and her healthy parents. Library preparation, sequencing (150 bp paired-end reads) and alignment for whole-exome sequencing were performed with the Genewiz exome-sequencing package. Potential disease-causing variants were investigated by Ingenuity Variant Analysis (Qiagen). Hight-quality variants were identified by filtering as follows: exclusion of common variants (>0.1% allele frequencies in public databases); retention of variants in coding regions resulting in substitutions, premature stops, frameshifts or altered splicing; exclusion of variants with CADD scores below MSC thresholds. For recessive models of inheritance, only homozygous or compound heterozygous mutations were assessed. For *de novo* inheritance, all variants in the parents were excluded, and alleles with known haploinsufficiency, hemizygous, or dominant-negative effects were included. For the validation of WES results, *JAK1* was then amplified from PBMC DNA by PCR and Sanger sequenced.

##### Digital droplet PCR

For the determination of mosaic allele fractions, DNA was isolated from bilateral buccal swabs, fractionated blood and a gastrointestinal endoscopic biopsy specimen in which the epithelial layer was isolated by chemical dissociation. Digital droplet PCR was performed with JAK1 primers and mutation-specific probes (IDT), and with ddPCR Supermix (Biorad 1863026). Amplification and quantification were performed on a QX100 Droplet Digital PCR system (BioRad). Cellular genotypes were estimated with QX100 software (BioRad), assuming heterozygosity.

##### Transduced- and patient-derived cell lines

The JAK1 plasmid was obtained from Addgene (Plasmid #23932) in a Gateway-compatible backbone, pDONR 223. Site-directed mutagenesis with specific primers (Quikchange II, Agilent 200521) was performed to obtain the WT CDS in the same plasmid, and plasmids encoding the S703I, K908A or S703I/K908A forms were then generated. A control vector containing the luciferase gene was also created. Plasmids were then subcloned into a lentivirus-compatible pTRIP-X-IRES-RFP backbone with puromycin resistance. Pseudotyped lentiviral particles were produced by the transfection of HEK293T cells with pCAGGS-VSV-G, pCMV-Gag/Pol and genes of interest. U4C cells (*JAK1*$^{-/-}$) were obtained from S. Pellegrini and transduced with lentiviruses at low MOI. Cells were selected on puromycin and FACS-sorted for matched RFP expression. EBV-transformed lymphoblastoid cell lines (EBV-B cells) were generated by infecting PBMCs from healthy controls or the patient with EBV. Single-cell clones were isolated by limiting dilution analysis on OP9 feeder cells and expanded in conditioned media. After genotyping by Sanger sequencing, WT/WT and WT/S703I clones were selected.

##### *In vitro* culture and stimulations

Transduced U4C cells were cultured in DMEM (Gibco) and EBV-B cells were cultured in RPMI, both supplemented with 10% fetal bovine serum (FBS) (Invitrogen), GlutaMAX (350 ng/ml; Gibco), and penicillin/streptomycin (Gibco). For the analysis of STAT phosphorylation, cells were stimulated with the indicated doses of recombinant IFNα-2b (Intron-A, Merck), IFNγ, IL-2, IL-4, and IL-6 (Biolegend) for 15 minutes and then lysed for western blotting. For the analysis of gene induction, cells were stimulated for 8 hours, after which cells were lysed for RNA isolation. For JAK inhibitor treatment, cells were incubated with ruxolitinib (Selleckchem S1378) or tofacitinib (Selleckchem S5001) at the indicated doses for 4 hours.

##### Immunoblotting

Cells were lysed in RIPA buffer (Thermo Fisher 89900) supplemented with protease/phosphatase inhibitor cocktail (Cell Signaling #5872). Lysates were sonicated, centrifuged to remove insoluble complexes, then boiled with NuPage sample buffer (Thermo Fisher NP0007) containing 20 mM DTT. The samples were subjected to gel electrophoresis and semi-dry transfer, and the resulting immunoblots were blocked in 5% BSA, then incubated overnight with primary antibody followed by HRP-conjugated secondary antibodies. Primary antibodies against the following targets were used: GAPDH (Cell Signaling D16H11), STAT1 (Santa Cruz C-111), phospho-STAT1 (Cell Signaling 58D6), phospho-STAT2 (Cell Signaling D3P2P), phospho-STAT3 (Cell Signaling D3A7), phospho-STAT5 (Cell Signaling C11C5), phospho-STAT6 (Cell Signaling 9361), JAK1 (Santa Cruz B3), JAK2 (Cell Signaling D2E12), TYK2 (Cell Signaling D4I5T), phospho-JAK1 (Cell Signaling 3331), phospho-JAK2 (Cell Signaling 3771), and phospho-TYK2 (Cell Signaling 9321). For the analysis of JAK phosphorylation, lysates were first incubated overnight with antibodies against total JAK protein conjugated to Protein G Dynabeads (Thermo Fisher 10007D). Immunoblotting was then performed as above.

**RT-qPCR**

Cell lines or isolated PBMCs were lysed and RNA was isolated with RNeasy spin columns (Qiagen 74104). Reverse transcription was performed with the High-Capacity RT Kit (Applied Biosystems 4368814). The resulting cDNA was then subjected to qPCR with the TaqMan Master Mix II with UNG (Thermo Fisher 4440038), on a Roche LightCycler 480, with the following primers/probes: *18S* (4318839), *MX1* (hs00895608), *RSAD2* (hs00369813), *SIGLEC1* (hs00988063) *IFIT1* (hs01911452). The relative expression of each transcript was normalized relative to *18S* by the ΔΔCt method.

**Mass cytometry**

Whole blood was collected, by venipuncture, into sodium heparin vacutainer tubes. For immunophenotyping, blood was immediately stained and processed for mass cytometry. For intracellular staining, whole blood was stimulated by incubation for 15 minutes with the cytokines indicated and was then immediately stabilized with Proteomic Stabilizer PROT1 (SmartTube) and frozen at -80°C. Similarly, for inhibition with ruxolitinib and tofacitinib, blood was treated for 4 hours with 500 nM inhibitor, and was then stimulated by incubation for 15 minutes with 1000 IU/mL IFNα, before stabilization and freezing.

Frozen samples were thawed according to the manufacturer's recommended protocol. The thawed samples were washed with barcode permeabilization buffer (Fluidigm) and barcoded with Fluidigm's Cell-ID 20-Plex Pd Barcoding Kit. Samples were then washed and pooled into a single tube, Fc-blocked (FcX, Biolegend) and heparin-blocked to prevent non-specific binding. Cells were then stained with a cocktail of markers to identify major immune populations. All antibodies in the panel were either conjugated in-house with X8 MaxPar conjugation kits (Fluidigm) or purchased from Fluidigm. The antibody cocktail was filtered through an Amicon filter with 0.1 µm pores before staining.

After surface staining, the samples were permeabilized with methanol and stored for at least 12 hours in methanol at -80°C. Samples were then washed, heparin-blocked and stained with a cocktail of phosphorylation and signaling antibodies. The stained samples were washed and incubated in freshly diluted 2.4% formaldehyde containing 125nM Ir Intercalator (Fluidigm), 0.02% saponin and 30 nM $OsO_4$ (ACROS Organics) for 30 minutes at room temperature. Samples were then washed and acquired immediately after staining.

Samples were washed once with PBS+0.2% BSA, once in PBS, and once in CAS buffer (Fluidigm). They were and resuspended at a concentration of 1 million cells per mL in CAS buffer containing a 1/20 dilution of EQ beads (Fluidigm). Following routine instrument tuning and optimization, samples were run at an acquisition rate of <300 events per second on a Helios mass cytometer (Fluidigm) with a modified wide-bore injector (Fluidigm). FCS files were then normalized and concatenated with Fluidigm acquisition software, and the barcoded samples were deconvoluted with a Matlab-based debarcoding application ("*Palladium-based mass tag cell barcoding with a doublet-filtering scheme and single-cell deconvolution algorithm*"). The FCS files were then uploaded to Cytobank for analysis. Cell events were identified as Ir191/193-positive and Ce140-negative events. Doublets were excluded on the basis of Mahalanobis distance and barcode separation and with the Gaussian parameters acquired with Helios CyTOF software. Downstream data analysis was performed on Cytobank, by both tSNE analysis and biaxial gating of immune populations, as shown in Supplementary Figure 3. Mean signal intensities were calculated, and relative induction was determined by normalization relative to the mean for healthy control samples.

**Flow cytometry**

NK cell phenotyping panels were modified from those described in Mahapatra et al (Mahapatra *et al.*, 2017). Cryopreserved PBMCs from the patient collected before and after the initiation of tofacitinib treatment, and from three unrelated healthy donors were thawed and allowed to rest briefly in complete RPMI medium supplemented with 10% FCS. Cells were immunostained with antibodies in 2% FBS in PBS for 45 minutes. For the panel assessing effector function and activation, cells were stimulated with phorbol 12-myristate 13-acetate (10 ng/ml, Sigma-Aldrich) and ionomycin (1 µg/ml, Sigma-Aldrich) for 4 hours at 37° C in the presence of brefeldin A (10 µg/ml, Sigma-Aldrich) and anti-CD107a antibody. For the panels evaluating effector function and transcription factors, cells were then permeabilized with BD Cytofix/Cytoperm (BD Biosciences) or FoxP3 buffer (Tonbo), and were stained by incubation with antibody for 45-60 minutes. Activated cells were stained for surface markers for 20-25 minutes after the four hours of incubation. Data were acquired on a FACSAria machine (BD Biosciences) with the capacity to detect 18 fluorescent parameters and exported to FlowJo 10.5.3 (TreeStar) for analysis. The frequency of cells positive for each parameter was compared with the mean and standard deviation for three healthy donors analyzed in parallel with the samples from the patient.

**Multiplex ELISA**
Plasma was collected by Ficoll isolation from heparinized whole blood, and clarified by centrifugation. Circulating cytokine levels were determined in magnetic Luminex assays with the Bio-Plex Pro Human Inflammation (BioRad 171al001m) and custom Human Cytokine Panel (R&D LXSAHM), according to the manufacturer's protocol. Samples were quantified on a MAGPIX xMAP Instrument (Luminex). Cytokine concentrations were quantified by comparison with standard curves and were subsequently translated into *Z*-scores.

**Immunohistochemistry**
Immunohistochemistry staining was performed with a Discovery Ultra instrument (Roche), with the staining module of RUO Discovery Multimer V2 (V0.00.0083). Slides were first incubated in blocking agent containing 2% BSA PBS, and Discovery Ultra antibody block (Cat # 760-4204) (Roche). Slides were incubated with primary antibodies for 60 minutes at 37°C. The following primary antibodies were used: anti-phospho-STAT1 (Tyr701) 58D6 (Cell Signaling) and anti-phospho-STAT3 (Tyr 705) D3A7 XP (Cell Signaling) at 1:100 dilution in blocking agent. The slides were then incubated with Omni Map anti-rabbit HRP-conjugated secondary antibody (Multimer HRP) (Cat # 760-4311) (Roche) for 32 minutes. Positive signals were detected with the Discovery ChromoMap DAB Kit (Cat #760-159) (Roche).

**Modified inDrop single cell RNA-Seq targeting JAK1 S703I site**
inDrop and related droplet microfluidics single cell RNA-Seq strategies coencapsulate single "barcode microbeads" and individual cells in droplets. Reverse transcription of polyadenylated (polyA) mRNA incorporates a cellular barcode sequence (different for each bead) into nascent cDNA. After downstream high throughput sequencing, reads can be assigned to individual cells by barcode sequences, thereby enabling expression quantification at single cell resolution. As the cellular barcode is introduced downstream of transcript polyA tails, sequence data is typically restricted to transcript regions immediately proximal to 3' termini. Because the S703I site is located at position 2402 from 5' transcript start and 2690 from the 3' transcript polyA tail, it is not accessible by standard droplet microfluidics single cell RNA-Seq platforms. Therefore, we adapted the inDrop method by generating custom barcode microbeads containing JAK1-specific primers (flanking the S703I site) in addition to polyT primer sequences, enabling more efficient JAK1 target capture and access to the S703I site. Following within-droplet reverse transcription, second strand synthesis and *in vitro* transcription amplification, samples are split into two parallel library preparations: one for standard polyT-primed libraries, and one for JAK1-targeted libraries. During data processing of resulting high throughput sequencing data, sequence reads from both libraries are assigned to individual cells based on shared barcode sequences. A detailed description of this approach appears below.

*JAK1-targeted hydrogel microbeads*
Barcoding hydrogel beads were prepared according to established inDrop protocol (Zilionis <i>et al.</i>, 2017)(Zilionis et al., 2017)(Zilionis et al., 2017)(Zilionis et al., 2017)(Zilionis et al., 2017)(Zilionis et al., 2017)(Zilionis et al., 2017)(Zilionis et al., 2017) with modified primers (Zilionis *et al.*, 2019) with the following modifications. For the second round of split-and-pool primer extension for barcode synthesis, hydrogel beads were added to microplate wells (n=384) containing oligonucleotide templates for both standard polyT primers and for an additional primer complementary to an S703I-adjacent region of the JAK1 transcript (11.53uM polyT oligonucleotide template, 2.3uM JAK1 oligonucleotide template) Within a given well, both polyT and JAK1 oligonucletide templates carried identical "barcode 2" sequences, ensuring that extended primers on hydrogels contained matching barcodes. Subsequent exonuclease processing steps included corresponding JAK1 primer complementary blocking oligonucleotides.

*JAK1-targeted library preparation*
Freshly isolated PBMC were co-encapsulated with JAK1-targeted hydrogel beads. The standard inDrop protocol was followed for reverse transcription, droplet breakage, second strand synthesis and in vitro transcription (IVT) amplification. IVT reactions (20 ul) were then split to two parallel library preparations: 10 ul of IVT product was

prepped for polyT-primed gene expression libraries according to the standard inDrop protocol, and 10 ul of IVT product (typically reserved as a "backup" aliquot) was prepped according to a JAK1-targeted protocol as follows. IVT products were reverse transcribed by SuperScript III with a JAK1-specific primer (with 5' extension containing Illumina adaptor sequence) flanking the S703I site (55C for 1 hr, 70C for 15 min). RT reactions were treated with RNase H (37C for 30 min, 65C for 20 min) to remove RNA template from cDNA heteroduplexes. Following purification on 1.5X Ampure XP beads (Beckman Coulter), JAK1-enriched cDNA was amplified by PCR with Illumina-adapted inDrop primers (KAPA HiFi Master Mix; 2 cycles 98C x 20 sec, 55C x 30 sec, 72C for 40 sec; 16-18 cycles 98C x 20 sec, 65C x 30 sec, 72C for 40 sec; final 72C extension x 5 min). Final JAK1-targeted libraries were purified on 0.8X Ampure XP beads.

*Oligonucleotide sequences*

polyT oligonucloeitde template 5'-BAAAAAAAAAAAAAAAAAAAANNNNNN [bc2, 8nt] CTGTCTCTTATACACATCTCCGAGCCCACG – 3'

JAK1 oligonucleotide template 5' - GAGTGTGGCCCATTCATCAANNNNNN [bc2, 8nt] CTGTCTCTTATACACATCTCCGAGCCCACG -3'

JAK1 blocking oligonucleotide 5'-GAGTGTGGCCCATTCATCAA-3'

Final "on bead" polyT primer sequence 5'-CGATGACGTAATACGACTCACTATAGGGTGTCGGGTGCAG[bc1,8nt]GTCTCGTGGGCTCGGAGATGTGTATAAGAGACAG[bc2,8nt]NNNNNNTTTTTTTTTTTTTTTTTTTV-3'

Final "on bead" JAK1 primer sequence 5'-CGATGACGTAATACGACTCACTATAGGGTGTCGGGTGCAG[bc1,8nt]GTCTCGTGGGCTCGGAGATGTGTATAAGAGACAG[bc2,8nt]NNNNNNTTGATGAATGGGCCACACTC-3;

Second reverse transcription JAK1 primer: 5'-TCGTCGGCAGCGTCAGATGTGTATAAGAGACAGCCATGGAAATTCAAAGTTGCCAAACAG-3'

*High Throughput Sequencing*

Both standard polyT-primed libraries and JAK1-targeted libraries were pooled together and sequenced in multiplex on the Illumina NextSeq 500 platform with 75-cycle reagent kits in paired-end, dual index configuration:

Read 1 containing transcript/JAK1 data: 61 cycles

i7 read containing cell barcode data: 8 cycles

i5 read containing sample index data: 8 cycles

Read 2 containing cell barcode and unique molecular identified (UMI) data: 14 cycles

*Data Processing*

FASTQ sequence files were processed with the indrops.py workflow script (v0.3, github.com/indrops/indrops). For polyT-primed gene expression libraries, resulting gene x cell matrices and per cell read counts were used for downstream analyses in Seurat (details below), using only cell-containing droplets with a read count higher than 8000-10000 depending on read count evaluation from droplets in the corresponding split library.

For JAK1-targeted libraries, BAM files (appended with cellular barcode and UMI data) were used to evaluate genotypes at single cell resolution as follows. Reads covering the JAK1 S703I site (chr1:64845519 – 64845521 on the minus strand; human genome reference GRCh38) containing the wildtype sequence (GAT) or S703I (GCT) sequence were quantified per UMI per cellular barcode. If at least 2 and at least 90% of the reads for a given cellular barcode/UMI combination contained the same sequence (wildtype or S703I), then this combination was designated a JAK1 transcript of the given genotype and assigned to the appropriate cellular barcode.

We next used per cell JAK1 WT and S703I transcript counts to assign putative genotypes to individual cells. As cell free RNA in suspension can co-encapsulate with cells thereby generating unwanted background signal in droplet microfluidics scRNA-Seq methods, we aimed to apply a stringent, data driven threshold for genotyping assignment. To evaluate the potential influence of cell free RNA, we quantified the frequency of JAK1 transcripts detected in empty (i.e. cell free) droplets (defined as barcodes with 1000 – 2000 reads in polyT-primed libraries) using the same barcode/UMI strategy described above. We found that more than 99% of empty droplets had less than 3 JAK1 transcripts. Guided by these data, cells were only considered for JAK1 genotype assignment if at least 5 JAK1 transcripts were detected. For JAK1 genotype assignment, individual cells were classified in one of the following four categories: cells that carried the S703I allele were classified as "MUT+" regardless of carrying the wildtype allele; the remainder (S703I allele negative) cells that carried the wildtype allele were classified as "WT+MUTneg" if no S703I transcripts were detected, and as "WT+MUTnonZero" if 1-to-4 S703I transcripts were detected; cells that carried neither the wildtype nor the S703I allele were classified as "WTnegMUTneg."

##### Single cell RNA-Seq Gene Expression Analysis

Data from JAK1-targeted inDrop single cell RNA-Seq data were analyzed in conjunction with corresponding single cell RNA-Seq data acquired from the same PBMC specimen on the 10X Genomics Chromium platform. Sequence reads from 10X libraries were processed with the CellRanger software package (10X Genomics) using default parameters. Gene x cell matrices from both methods were further analyzed with Seurat (v2..3.4) (Butler *et al.*, 2018) in the R statistical framework as follows. For applicable cells in the JAK1-targeted inDrop dataset, JAK1 genotyping information was imported as per cell metadata entries. Genes with detectable expression in fewer than 5 cells per sample were excluded. Cells with detectable expression of fewer than 250 genes or fewer than 1000 UMI counts or greater than 20% UMIs from mitochondrial gene transcripts (measure of cell viability) were removed from further analysis. Gene expression data were log normalized and scaled (regressing out effects based on total UMI counts and mitochondrial gene expression). All genes with detectable expression in both inDrop and 10X datasets were used for CCA (20 dimensions). CCA dimension scores were aligned between datasets using the AlignSubspace function. Aligned data were log-normalized and scaled as above. Graph-based clustering and tSNE visualization was performed with the FindClusters and RunTSNE functions with default settings (using the first 19 CCA dimensions, based on examination of distinct expression patterns with the DimHeatmap function). Genes distinguishing cell clusters (n = 14) were identified with the FindMarkers function. Cell type clusters were manually annotated based on marker gene expression patterns.

Gene expression differences were assessed for cells assigned JAK1 genotype classifications of either MUT+ (S703I) or WT+MUTneg (WT). Plasmablasts and dendritic cells were excluded from analysis due to insufficient cells assigned to both genotype groups. Apparent cellular doublets and cells in the platelet cluster were also excluded. Differential gene expression testing was performed on genes detected in at least 20% of cells in any cell type cluster (of either JAK1 genoytpe group) with edgeR (v.3.12.1)(Robinson, Mccarthy and Smyth, 2010), including modifications included for single cell RNA-Seq data as described (Soneson and Robinson, 2018). A linear model including factors for cellular gene detection rate, cell type, JAK1 genotype group and an interaction term (cell type cluster and JAK1 genotype group), was fit with the glmQLFit function. Differential gene expression testing (WT vs. S703I) was tested across all cell types with the glmQLFtest function, with significance thresholds set at FDR 0.05. In addition, gene set enrichment testing for Molecular Signatures Database Collections H, C2-CPC, C3-TFT and C7 collections (http://software.broadinstitute.org/gsea/msigdb), supplemented with two additional interferon stimulated gene sets (Rosenberg *et al.*, 2018) was performed with CAMERA (Wu and Smyth, 2012) and the linear model described above. Only gene sets with 5 or more genes in the fitted model were included in the analysis.

##### Single Cell qPCR Transcript Genotyping

PBMCs from a healthy donor carrying a heterozygous SNP in *JAK1 (*rs2230587*)* were isolated by Ficoll gradient. Cells were stained with antibodies against CD3, CD19, CD14 and CD56 (Biolegend), and 100 single cells were FACS-sorted into single cell lysis buffer (Ambion 4458235). Following DNAse treatment, cDNA was generated using SuperScript VILO RT kit (ThermoFisher 11754050). A linear preamplification was then performed using *JAK1* primers (gtcctctggatctcttcatgca, gctgtttggcaactttgaatttcc) and primers to a negative control gene *NACA* (cccaggcaaccacacaac, ccgactctgttttgctttactgact). Using qPCR with custom TaqMan genotyping primers (above) and allele-specific probes (*JAK1* aaggacatc[g/a]cttttc; agcagctgaaat[T/C]gatgaa) that were individually fluorescently tagged (VIC and FAM), allelic ratios were determined by endpoint genotyping. Quantification was carried out by interpolation from a standard curve of oligonucleotide standards.
